# nnOPN3 mediates retinal-dependent lipofuscin accumulation and its loss sensitizes keratinocytes to blue-light-induced proteomic remodeling

**DOI:** 10.64898/2026.09.09.750435

**Authors:** Maiza Von Dentz, Helena Couto Junqueira, Manuel Alejandro Herrera Lopez, Gabriella Lisboa, Johannes Fuchs, Katherine Tsantarlis, Carina Sihlbom Wallem, Leonardo Vinicius Monteiro de Assis, Maurício da Silva Baptista

## Abstract

Lipofuscin is a blue-light-absorbing pigment that contributes to oxidative damage. Whether *all-trans* retinal (atRAL) contributes to its formation in response to light remains unclear. We asked whether blue-light photosensitization of atRAL promotes lipofuscin accumulation in human keratinocytes and whether this depends on Opsin 3 (OPN3). Blue-light excitation of atRAL reduced mitochondrial and lysosomal viability, impaired autophagic flux, and increased lipofuscin. OPN3 knockdown significantly reduced this accumulation. Label-free data-independent acquisition (DIA) proteomics showed that OPN3 acts at three levels. In the dark, OPN3 loss altered proteome networks associated with autophagy, apoptosis, and interferon-related responses. Under blue light, control cells activated a stress-adaptive program spanning inflammatory regulation, lipid metabolism, and mitochondrial function, which atRAL strongly amplified. This molecular signature, including induction of cellular respiration and ATP-production proteins, was largely absent when OPN3 was silenced. Respirometry showed that blue light suppressed oxygen consumption in both lines over the first 24 h, but a faster recovery in OPN3 knockdown cells at 48–72 h was observed. Together, these data define three functions of OPN3: maintaining the basal proteome in a blue-light-independent manner, enabling the adaptive blue-light response, and enabling retinal-dependent lipofuscin formation in keratinocytes.

## INTRODUCTION

Visible light, especially in the blue-light range, has gained attention as an important source of oxidative and metabolic stress in skin cells ^1–3^. Increasing evidence shows that visible light can induce reactive oxygen species, inflammatory mediators, and matrix-degrading enzymes in many skin models ^4,5^. One important aspect is that blue light can interact with endogenous photosensitizers naturally present in cells, including flavins, porphyrins, melanin, heme-containing molecules, light sensors (opsins) and lipofuscin, triggering the generation of reactive species and secondary signaling pathways ^4^.

Lipofuscin is a potent photosensitizer that accumulates in aging cells and absorbs blue light and generates reactive species that can trigger mutagenic lesions ^6–8^. This is exemplified by its role in generating mutagenic DNA lesions in keratinocytes through singlet oxygen (¹O₂)-mediated pathways upon visible light exposure ^9^. Lipofuscin is generally described as being composed of oxidized proteins, lipids, carbohydrates, and metals such as iron and calcium ^10–12^. Cells do not have efficient machinery to process lipofuscin, so this pigment accumulates over time in tissues, creating a vicious cycle of production and damage ^13,14^. Mechanisms of lipofuscin generation include inhibition of autophagic flux through damage to the lysosomal–mitochondrial axis and, at least in retinal pigment epithelial cells, imbalance of the retinoid visual cycle ^15,16^. It has been shown in keratinocytes that inhibiting autophagy can lead to the buildup of lipofuscin ^8^. In retinal pigment epithelial cells, lipofuscin formation has been linked to the visual cycle, where A2E (N-retinylidene-N-retinylethanolamine), a dimer of retinaldehyde, is generated as a byproduct ^17^. However, whether the monomeric form of retinal (*all-trans* retinal, atRAL) itself can act as a photosensitizer to induce lipofuscin formation has not been investigated. This is a critical gap because atRAL is present not only in the retina but also in keratinocytes and other peripheral cells, where it is derived from vitamin A metabolism ^18^.

Opsins are G-protein coupled receptors that bind covalently to molecules of 11-cis retinal. After absorption of light, 11-cis retinal is isomerized into atRAL and delivered to the cell. Once in the cell, atRAL can be regenerated or contribute to lipofuscin components ^17,19^. Among the opsins, OPN3 is highly expressed in skin cells and has been associated with pigmentation, apoptosis, proliferation, metalloproteinase activity, and blue-light responses ^20–23^. Recently, OPN3 photoactivation by blue light was also related to autophagy inhibition in melanocytes through an OPN3–TRPV1–calcium influx pathway ^24^. However, the relationship between OPN3, retinal photosensitization, and lipofuscin formation remains unexplored.

We investigated whether *all-trans* retinal acts as an endogenous photosensitizer capable of promoting lipofuscin accumulation in human keratinocytes after blue-light exposure, and whether this response depends on OPN3. Our study revealed a complex function of OPN3, displaying both light-dependent and light-independent roles associated with retinal and lipofuscin accumulation.

## RESULTS

### Blue light photosensitizes atRAL and induces lipofuscin accumulation

atRAL is an endogenous molecule derived from the metabolism of vitamin A. It is an oxidized form of retinol, produced in cells by retinol dehydrogenases in an NADP-dependent manner ^25^. While retinoic acid and its derivatives, such as retinol and retinal, are considered beneficial for anti-aging effects, their use is typically restricted to nighttime due to their photoinstability and association with photosensitivity and skin irritation ^26,27^. Moreover, even without external formulations, physiological serum concentrations of retinol can reach up to 2.5 µM, allowing cells to take up and convert it into retinal isoforms ^28,29^. Although the maximum absorption of atRAL is at 383 nm ^30,31^, we show that blue light acts as a photosensitizer for atRAL.

We hypothesized that atRAL photosensitization could be related to an imbalance in the retinoid visual cycle and autophagic flux. To test this hypothesis, we conducted experiments to measure lipofuscin accumulation and assess autophagic flux integrity. Lipofuscin content was measured through autofluorescence (Fig. 1). While atRAL itself does not cause an increase in lipofuscin content, exposure to blue light after atRAL incubation raised it by approximately 2.5-fold (Fig. 1 D; E). In contrast, blue light alone does not affect this parameter compared to the dark group (Fig. 1 C; E). This is novel evidence showing that photosensitization of atRAL, rather than its conversion to A2E or other bisretinoids, can drive lipofuscin accumulation. The increase in lipofuscin in skin cells has also been associated with chronic UVA exposure or parallel damage of mitochondrial and lysosomal function using photosensitization of DMMB with red light ^8,9^. In our experimental conditions, the increase in lipofuscin was associated with decreased mitochondrial viability and membrane integrity (Fig. S1), suggesting that atRAL photosensitization initiates a cascade of organelle damage.

**Fig. 1.**
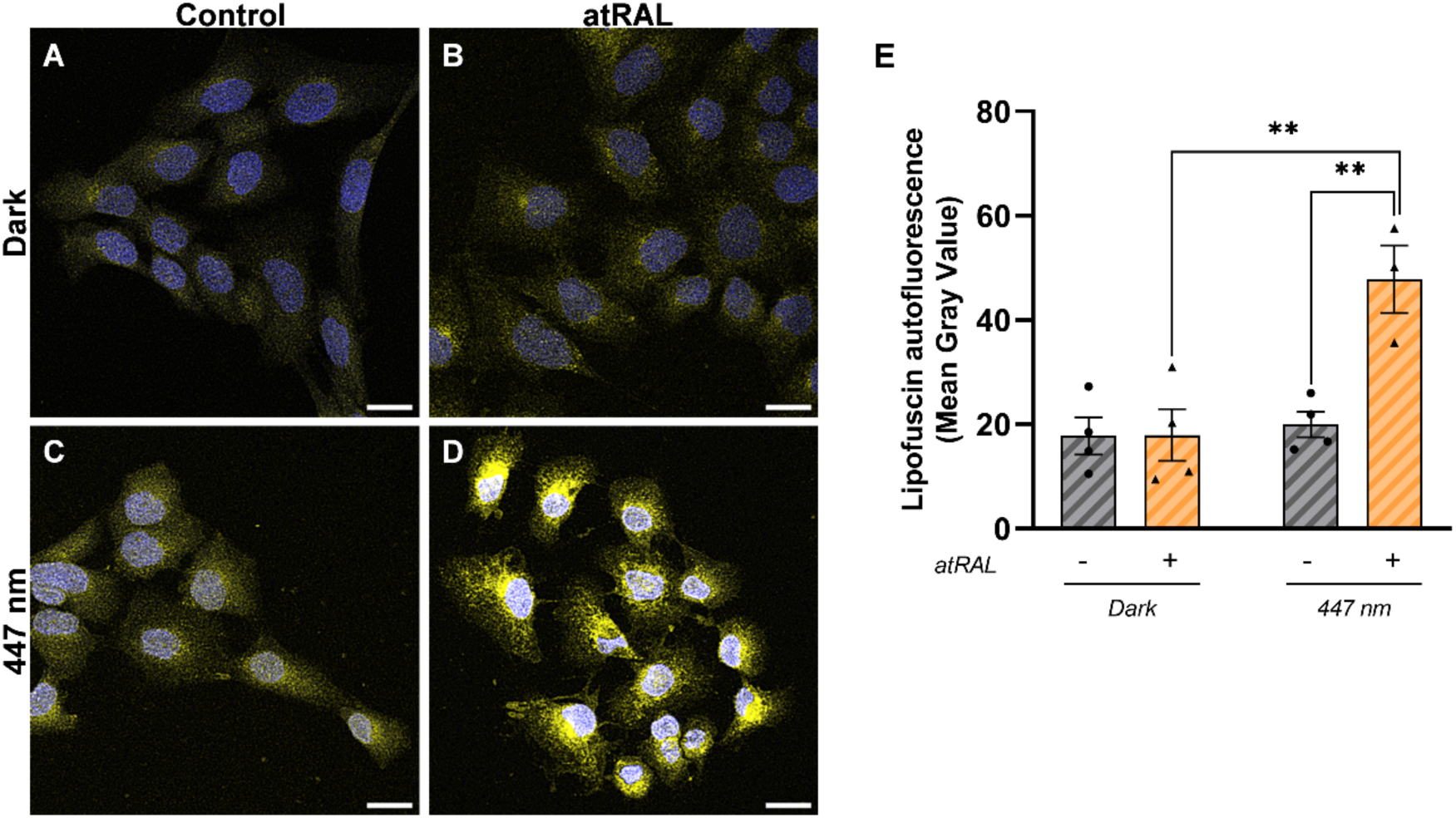
Assessment of lipofuscin accumulation 48 h after atRAL photosensitization. (A-D) Representative images of lipofuscin autofluorescence (Ex. 488 nm/Em. 509–601 nm). (E) Quantification of lipofuscin autofluorescence as mean gray value. Experimental groups were: (A) Dark Control, (B) Dark + atRAL, (C) 447 nm Control, and (D) 447 nm + atRAL. A total of 50 cells per group were analyzed for autofluorescence quantification. Each dot in the graphs represents one independent biological replicate, with measurements obtained from the indicated number of cells per replicate. Data are presented as the mean ± SEM from at least three independent experiments. Statistical analysis was performed using two-way ANOVA followed by Bonferroni’s multiple-comparisons test. ANOVA effects: interaction (*P* = 0.0088), irradiation (*P* = 0.0036) and retinal treatment (*P* = 0.0083). \**P* < 0.05.

### atRAL photosensitization inhibits autophagic flux and increases acidic vacuole accumulation

An established way to accumulate lipofuscin in cells is through blocking autophagic flux ^15^. To confirm whether the autophagic flux is affected by atRAL photosensitization, we employed viability results in a mathematical formula that showed that photosensitization of atRAL increases inhibition of autophagic flux 48 h after treatments (Fig. 2 A) ^32^. The same data were obtained by LC3-II net flux and steady-state assays performed 24 hours after treatments (Fig. 2 B - D). These assays use bafilomycin to inhibit autophagic flux in each experimental condition and use the differences between groups with and without bafilomycin to determine whether increased levels of LC3-II reflect an increase or decrease in autophagic flux, which is considered one of the best ways to evaluate autophagy ^33,34^. Our findings show that atRAL plus blue light has an additional effect on the inhibition of autophagy in relation to blue light treatment alone, as indicated by steady-state LC3-II levels (Fig. 2 B, D) and LC3AB-II net flux (Fig. 2 C).

**Fig. 2.**
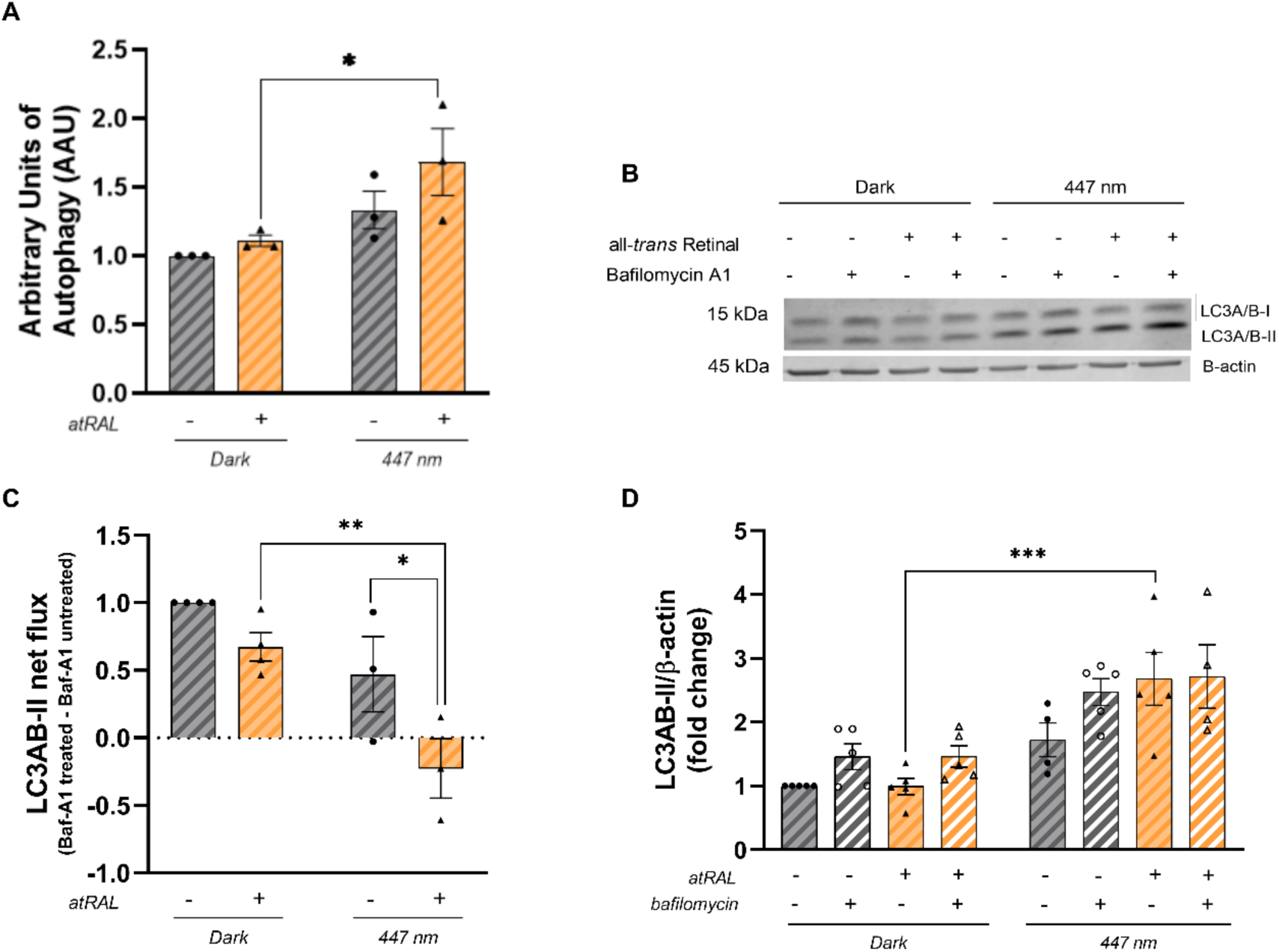
Assessment of autophagic flux following atRAL photosensitization. (A) Quantification of autophagic flux expressed as autophagy arbitrary units (AAU) measured 48 h after treatment. (B) Representative immunoblot of LC3-II and β-actin 24 h after treatment. For each experimental condition, paired samples were treated with 200 nM bafilomycin A1 for 1 h before protein extraction to assess autophagic flux. (C) Quantification of LC3-II net flux. (D) Quantification of LC3-II steady-state levels (fold change). Experimental groups were Dark Control, Dark + atRAL, 447 nm Control, and 447 nm + atRAL. Each dot in the graphs represents one independent biological replicate. Data are presented as the mean ± SEM from at least three independent experiments. Statistical analysis was performed using two-way ANOVA followed by Bonferroni’s multiple-comparisons test for panels A and C, and three-way ANOVA followed by Bonferroni’s multiple-comparisons test for panel D. ANOVA effects: AAU— interaction (*P* = 0.4173); irradiation (*P* = 0.0120); retinal treatment (*P* = 0.1398). LC3-II net flux—interaction (*P* = 0.2739); irradiation (*P* = 0.0012); retinal treatment (*P* = 0.0096). LC3-II steady-state levels—three-way interaction (*P* = 0.3416); irradiation (*P* < 0.0001); retinal treatment (*P* = 0.1194); bafilomycin treatment (*P* = 0.0290); irradiation × retinal treatment (*P* = 0.1178); irradiation × bafilomycin treatment (*P* = 0.8450); retinal treatment × bafilomycin treatment (*P* = 0.3567). \**P* < 0.05.

Labeling trackers of acid vacuoles is also used to evaluate lysosomal function. Three hours and 24 hours after treatments, cells were labeled with lysotracker deep red or acridine orange. No changes in fluorescence were detected 3 hours after treatment (Fig. S2), but 24 hours after, atRAL photosensitization increased fluorescence in cells labeled with lysotracker (Fig. 3 A – E) and acridine orange (Fig. 3 F – J). There were no changes in the immunocontent of LAMP1 (Fig. S3), a lysosomal membrane protein, suggesting that increasing fluorescence is related to autophagosome accumulation and downstream inhibition of autophagic flux ^35,36^.

**Fig. 3.**
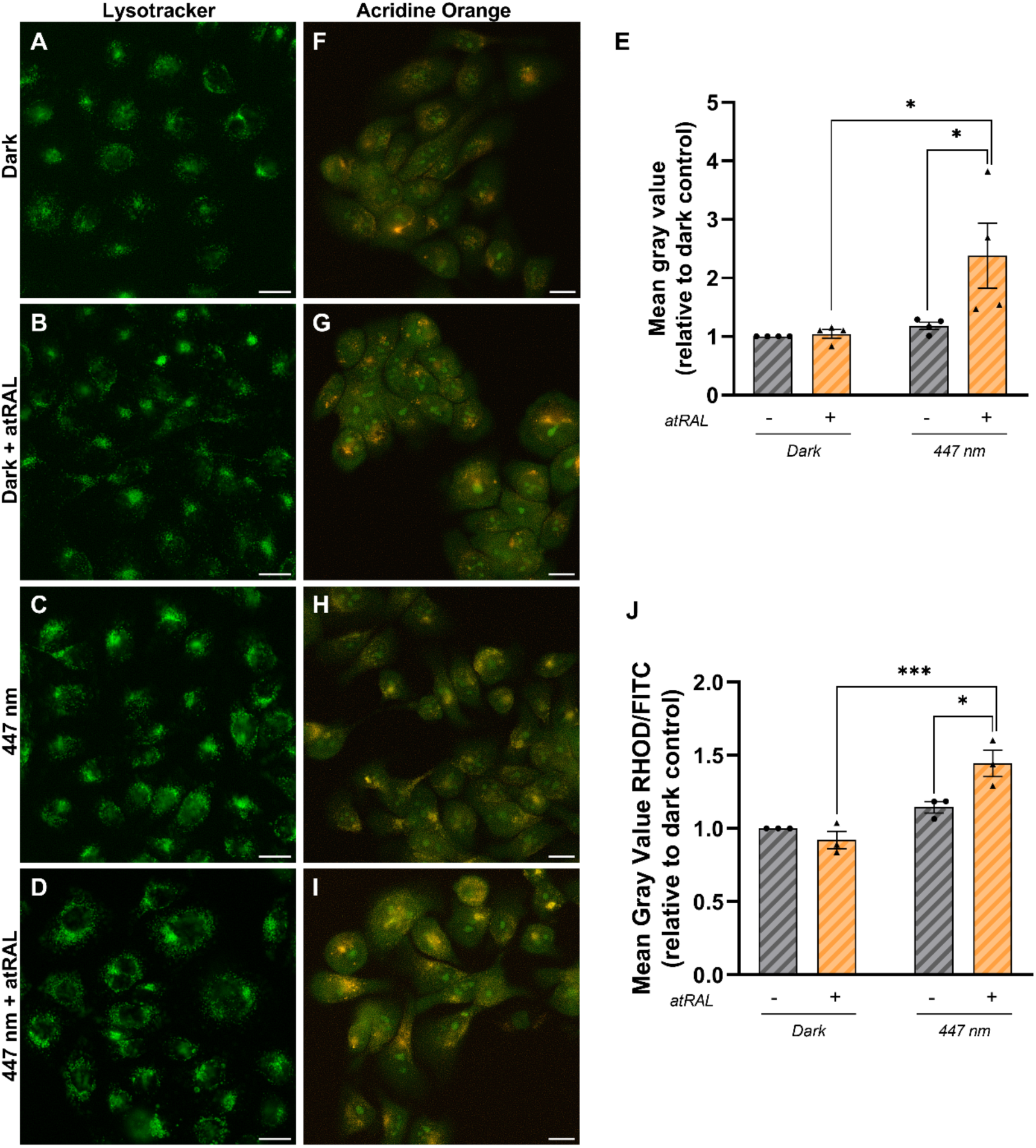
Assessment of acidic vacuoles 24 h after atRAL photosensitization. (A–D) Representative images of LysoTracker Deep Red staining in the different experimental groups. (E) Quantification of LysoTracker Deep Red fluorescence as mean gray value normalized to the Dark Control group. (F–I) Representative images of acridine orange fluorescence acquired in the FITC and Rhodamine (RHOD) channels. (J) Quantification of the acridine orange RHOD/FITC fluorescence ratio as mean gray value normalized to the Dark Control group. Experimental groups were: (A,F) Dark Control, (B,G) Dark + atRAL, (C,H) 447 nm Control, and (D,I) 447 nm + atRAL. A total of 50 cells per group were analyzed for each assay. Each dot in the graphs represents one independent biological replicate, with measurements obtained from the indicated number of cells per replicate. Data are presented as the mean ± SEM from at least three independent experiments. Statistical analysis was performed using two-way ANOVA followed by Bonferroni’s multiple-comparisons test. ANOVA effects: LysoTracker Deep Red— interaction (*P* = 0.0630); irradiation (*P* = 0.0198); retinal treatment (*P* = 0.0478). Acridine orange RHOD/FITC ratio—interaction (*P* = 0.0113); irradiation (*P* = 0.0004); retinal treatment (*P* = 0.0912). \**P* < 0.05.

Although blue light irradiation alone increased inhibition of autophagic flux (Fig. 2 B – D), only atRAL photosensitization was sufficient to increase acid vacuole fluorescence (Fig. 3A – J). Together, these results showed that atRAL photosensitization promotes long-term inhibition of autophagic flux at downstream steps, including inhibition of autophagosome-lysosome fusion, which can trigger lipofuscin accumulation.

### OPN3 downregulation abolishes the lipofuscin increase induced by atRAL photosensitization

The inhibition of autophagic flux partially explains how atRAL photosensitization promotes an increase in lipofuscin, although the specific mechanistic process remains unclear. In the retinoid visual cycle, retinal and blue light are closely associated with OPN3, a GPCR highly expressed in keratinocytes. A recent study has described blue-light activation of the OPN3–TRPV1–calcium influx pathway to inhibition of autophagy in melanocytes ^24^. In our experimental model, atRAL photosensitization by blue light increased *OPN3* expression (Fig. S4). Based on these findings, we hypothesized that atRAL photosensitization under blue light could require OPN3 to regulate autophagic flux and promote lipofuscin accumulation.

To address this question, we generated cells deficient in OPN3, which expressed only 15% of *OPN3* compared to cells with the scrambled sequence (shCR) (Fig. 4A). Both shCR and shOpn3 cells were subjected to identical photosensitization protocols, and lipofuscin content was measured via autofluorescence 48 hours after treatment. While non-irradiated cells responded similarly to atRAL incubation regardless of OPN3 levels, with shCR controls showing a slight increase in lipofuscin accumulation (Fig. S5), the atRAL photosensitization effect on lipofuscin accumulation was strongly attenuated when OPN3 was downregulated (Fig. 4B–F). Notably, this effect was not due to reduced atRAL uptake, as intracellular and extracellular atRAL levels were similar between shCR and shOpn3 cells (Fig. S6). These findings reveal that OPN3 is required for atRAL-dependent lipofuscin formation through a mechanism independent of retinal transport or uptake.

**Fig. 4.**
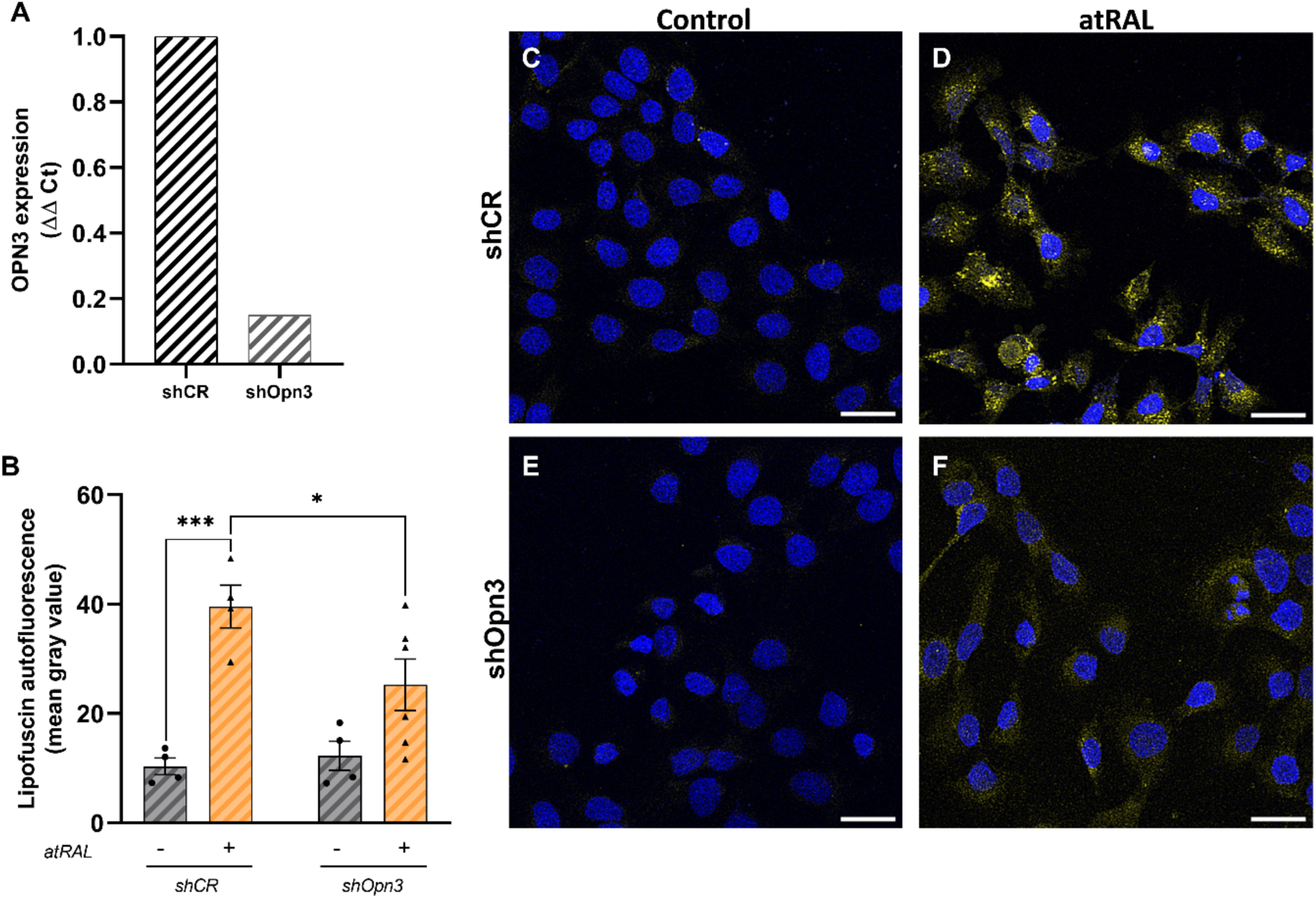
Assessment of lipofuscin accumulation in OPN3-knockdown cells following atRAL photosensitization. (A) Relative OPN3 expression in shCR and shOpn3 cells. (B) Quantification of lipofuscin autofluorescence as mean gray value. (C–F) Representative confocal images of lipofuscin autofluorescence (Ex. 488 nm/Em. 509–601 nm). Experimental groups were: (C) 447 nm shCR Control, (D) 447 nm shCR + atRAL, (E) 447 nm shOpn3 Control, and (F) 447 nm shOpn3 + atRAL. A total of 50 cells per group were analyzed for lipofuscin autofluorescence. Each dot in the graphs represents one independent biological replicate, with measurements obtained from the indicated number of cells per replicate. Data are presented as the mean ± SEM from at least three independent experiments. Statistical analysis was performed using two-way ANOVA followed by Bonferroni’s multiple-comparisons test. ANOVA effects: Lipofuscin autofluorescence (B)—interaction (*P* = 0.0594); OPN3-knockdown (*P* = 0.1392); retinal treatment (*P* = 0.0001). \**P* < 0.05.

### Basal OPN3-dependent proteomic remodeling is modestly affected by atRAL in the absence of light

To explore OPN3’s overall role in atRAL photosensitization, we conducted label-free proteomics on shCR and shOpn3 cells kept in darkness or exposed to blue light, with or without atRAL. Bioinformatic analysis using limma involved pairwise comparisons to assess each experimental factor’s impact. These analyses revealed a diverse range of effects across various comparisons (Fig. 5A; Table 1).

**Fig. 5:**
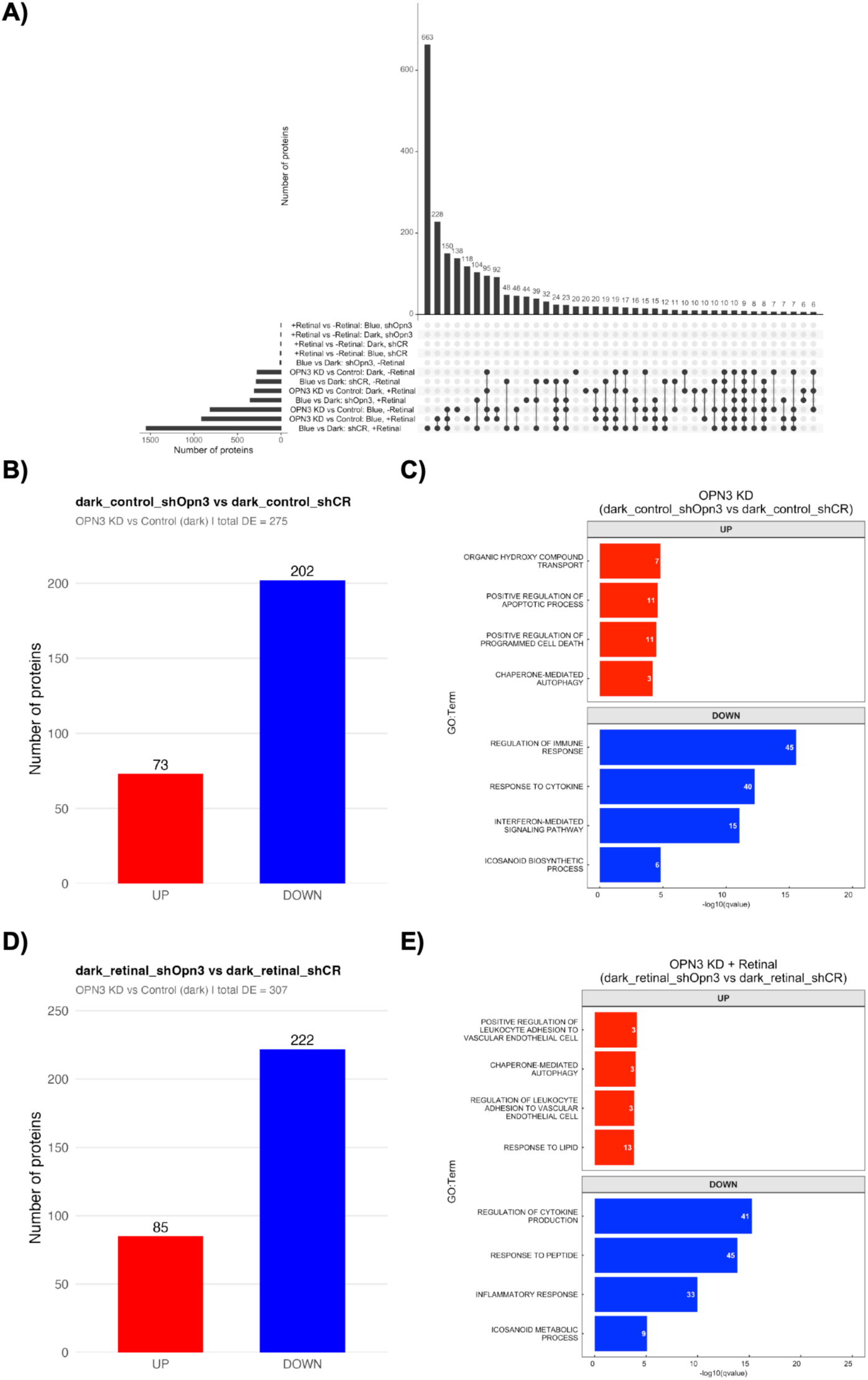
The cellular proteome is markedly affected by OPN3 silencing, blue light, and atRAL photosensitization. A) UpSet shows interactions among DEPs within each group. B and D) Histograms show the numbers of UP- and DOWN-regulated DEPs. C and E) Horizontal bar plots show the enriched biological processes according to identified DEPs.

**Table 1:** Overview of the identified differentially expressed proteins (DEPs).

| DEPs | Comparison | Description |
| --- | --- | --- |
| 275 | dark ctrl: shOpn3 vs shCR | Effect of OPN3 knockdown in dark conditions without atRAL |
| 307 | dark ret: shOpn3 vs shCR | Effect of OPN3 knockdown in dark conditions with atRAL |
| 813 | blue ctrl: shOpn3 vs shCR | Effect of OPN3 knockdown in blue light without atRAL |
| 912 | blue ret: shOpn3 vs shCR | Effect of OPN3 knockdown in blue light with atRAL |
| 5 | dark: retinal vs ctrl – shCR | Effect of atRAL supplementation in dark conditions in the shCR background |
| 9 | blue: retinal vs ctrl – shCR | Effect of atRAL supplementation in blue light in the shCR background |
| 4 | dark: retinal vs ctrl – shOpn3 | Effect of atRAL supplementation in dark conditions in the OPN3-knockdown background |
| 2 | blue: retinal vs ctrl – shOpn3 | Effect of atRAL supplementation in blue light in the OPN3-knockdown background |
| 284 | blue vs dark ctrl – shCR | Effect of blue light exposure without atRAL in the shCR background |
| 1547 | blue vs dark retinal – shCR | Effect of blue light exposure with atRAL in the shCR background |
| 12 | blue vs dark ctrl – shOpn3 | Effect of blue light exposure without atRAL in the OPN3-knockdown background |
| 354 | blue vs dark retinal – shOpn3 | Effect of blue light exposure with atRAL in the OPN3-knockdown background |

We identified several DEPs associated with a basal, i.e., light-independent, role of OPN3 in regulating critical biological processes. For instance, a total of 73 and 202 DEPs were up- and downregulated, respectively (BH < 0.05 & log2 fold change ± 0.58), in unexposed cells with reduced OPN3 expression compared with shCR cells (Fig. 5B; Table S1). Enrichment of the upregulated DEPs indicated that OPN3 silencing under basal conditions was associated with biological processes related to organic hydroxy compound transport (NOS1, NFKBIA, CLU), positive regulation of apoptotic signaling (DAPK1, RARG, CDKN2A), and chaperone-mediated autophagy (EEF1A2, CLU, SNCA). In contrast, downregulated DEPs were strongly enriched in immune- and inflammatory-related biological processes, including defense response to virus (OAS1, IFIT1, MX1), response to cytokines (STAT1, IRF7, ISG15), regulation of immune response (DDX58/RIG-I, IFIH1, USP18), antigen processing and presentation (HLA-A, TAP1, TAP2), and regulation of cytokine production (IL1B, CASP1, TNFAIP3) (Fig. 5 C; Table S3).

When atRAL was present in the dark in OPN3-silenced cells, the proteomic profile was largely similar to the basal OPN3-silencing condition. In this comparison, 307 DEPs were identified in shOpn3 cells compared with shCR cells (Fig. 5D, Table S1). Among the upregulated DEPs, the main enriched processes were related to regulation of leukocyte adhesion to vascular endothelial cells (ITGA4, MDK, IRAK1), chaperone-mediated autophagy (EEF1A2, CLU, SNCA), and response to lipid (RARG, SCD, SREBF1) (Fig. 5E; Table S3). Consistent with the basal OPN3-silencing profile, downregulated DEPs were mainly enriched in immune- and inflammatory-related biological processes, including defense response to virus (OAS2, IFIT1, MX1), response to cytokines (STAT1, IRF7, ISG15), regulation of innate immune response (DDX58/RIG-I, IFIH1, USP18), regulation of cytokine production (IL1B, CASP1, TNFAIP3), and antigen processing and presentation (HLA-B, TAP1, TAP2) (Fig. 5E; Table S3). Analysis of the exclusive DEPs further confirmed that atRAL under dark conditions has a minor impact. Only a small group of proteins was specific to the dark atRAL OPN3 comparison, totaling 20 DEPs (see Table S2). The enrichment analysis of these exclusive DEPs showed only a few affected biological processes, indicating that the molecular signature is largely similar regardless of whether OPN3 is knocked down in the presence or absence of atRAL (Table S4).

These findings suggest OPN3 helps maintain basal proteome homeostasis independently of light and that atRAL has a modest effect on the OPN3-dependent proteomic profile in the dark.

### atRAL amplifies the blue-light proteomic response and reveals a metabolic remodeling signature

We next evaluated the effect of blue light in shCR cells in the absence or presence of atRAL. In control cells without atRAL supplementation, blue light induced a moderate proteomic response, with 284 DEPs, including 245 upregulated and 39 downregulated proteins (Fig. 6A; Table S1). Enrichment analysis showed that blue light alone mainly increased biological processes associated with regulation of cell migration (BMPR1A, MMP14, APP), response to wounding (ATP7A, GRN, APOE), high-density lipoprotein particle clearance (LDLR, APOE, APOA1), and regulation of cell adhesion (MMP14, TFRC, TGFBR2) (Fig. 6B; Table S3). In contrast, downregulated proteins were enriched in immune- and interferon-related processes, including cellular response to type I interferon (USP18, IFIT1, IRF7), type I interferon production (IRF7, IFIH1, RIGI), response to cytokine (USP18, IFIT1, ISG15), and amino acid biosynthetic process (SRR, DPYD, CBS).

**Fig. 6:**
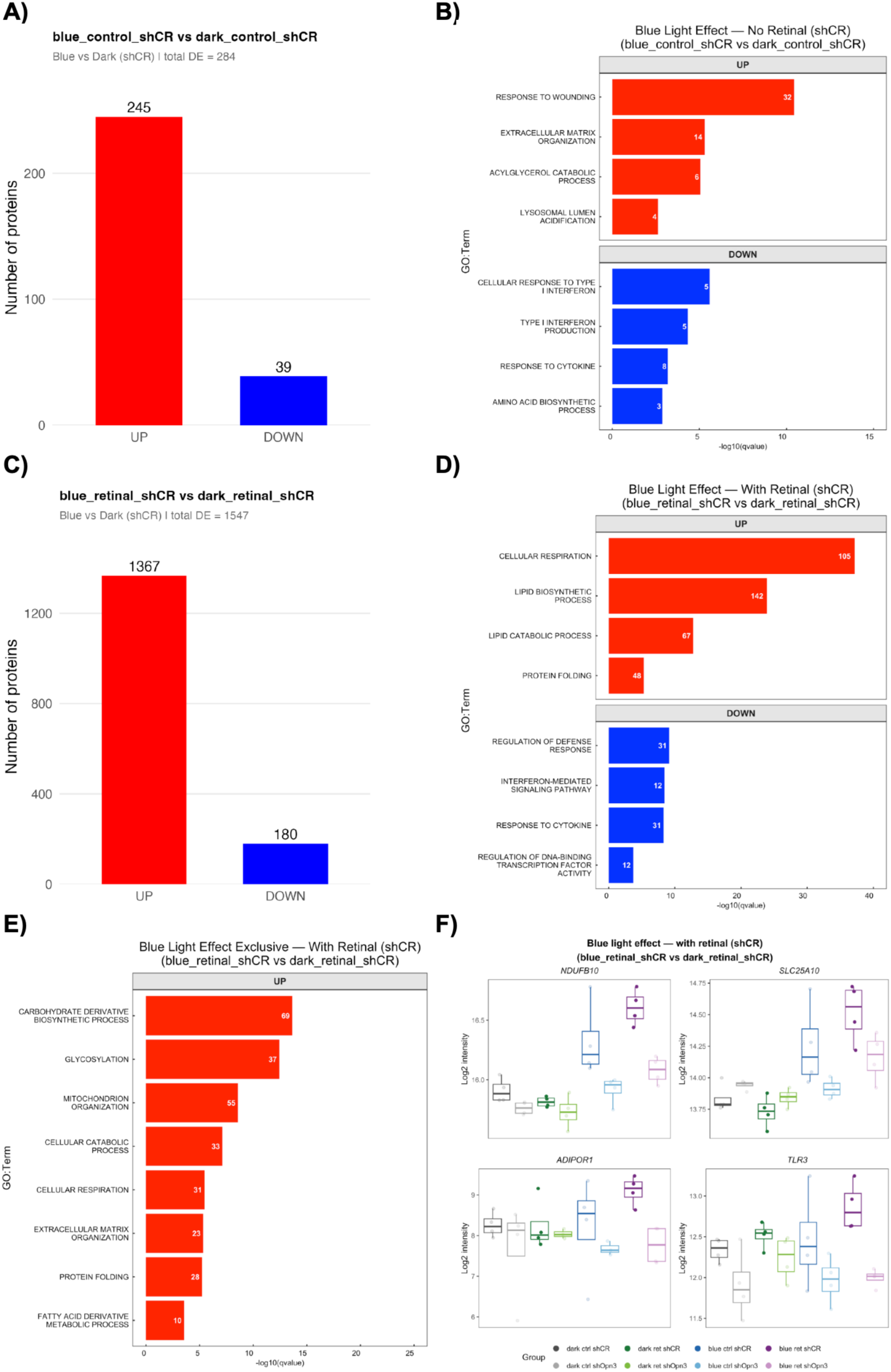
Blue-induced retinal photosensitization leads to a specific molecular signature. A and C) Histograms show the numbers of UP- and DOWN-regulated DEPs. B and D) Horizontal bar plots show the enriched biological processes according to identified DEPs. E) Exclusive biological processes identified in response to retinal photosensitization. F) Representative DEPs from the identified processes in E.

In contrast, when atRAL was present, blue light produced a markedly stronger proteomic response. A total of 1547 DEPs were identified in blue-light-exposed atRAL-treated cells compared with dark atRAL-treated cells, including 1367 upregulated and 180 downregulated proteins (Fig. 6C; Table S1). The 1367 upregulated proteins revealed a strong metabolic and organelle-associated signature, with enrichment of carbohydrate derivative biosynthetic process (DOLK, SLC35B2, B3GALT6), glycoprotein metabolic process (DOLK, MGAT2, MAN2A1), cellular respiration (NDUFB11, ATP5F1C, NDUFS8), oxidative phosphorylation (NDUFB11, ATP5F1C, COX4I1), monoatomic ion transmembrane transport (ATP6V0A2, CLCN7, MCOLN1), and lipid biosynthetic process (SPTLC1, KDSR, HSD17B12) (Fig. 6D; Table S3). The downregulated DEPs in atRAL-treated blue-light-exposed cells were enriched for immune-related processes, including response to type I interferon (IFIT1, STAT1, OAS1), interferon-mediated signaling pathway (USP18, STAT1, IRF1), interleukin-27-mediated signaling pathway (STAT1, OAS1, MX1), and type I interferon-mediated signaling pathway (USP18, STAT1, IFIH1) (Fig. 6D; Table S3).

To identify the atRAL-dependent component of the blue-light response, we next analyzed DEPs exclusive to blue-light exposure in the presence of retinal. This analysis identified 663 exclusive DEPs, including 605 upregulated and 58 downregulated proteins (Table S2). Exclusive GO enrichment was detected only among the upregulated proteins. The most relevant enriched metabolic processes included glycoprotein metabolic process (DOLK, B3GALT6, MGAT2), carbohydrate derivative biosynthetic process (DOLK, SLC35B2, B3GALT6), protein glycosylation (MGAT2, MAN2A1, ALG2), lipid biosynthetic process (SPTLC1, KDSR, HSD17B12), phospholipid metabolic process (PTDSS2, AGPAT2, LPCAT1), ceramide and sphingolipid biosynthetic processes (SPTLC1, KDSR, ELOVL1), glycerolipid metabolic process (SACM1L, PTDSS2, AGPAT2), fatty acid catabolic process and fatty acid oxidation (ABCD3, CPT1A, ECI1), steroid and cholesterol biosynthetic processes (HSD17B12, LSS, SREBF2), and cellular respiration/oxidative phosphorylation (NDUFB10, NDUFAF2, UQCC2) (Fig. 6E; Table S4). Additional processes also included enrichment of mitochondrial respiratory chain complex assembly (NDUFB10, NDUFAF2, UQCC2), extracellular matrix organization (DAG1, ITGA2, LAMB2), and response to protein folding (HSPD1, EIF2AK3, EDEM2). Representative proteins from this exclusive signature, including NDUFB10, SLC25A10, ADIPOR1, and TLR3, showed increased abundance specifically in the blue-light plus atRAL condition (Fig. 6F).

These findings show that the presence of atRAL does not simply intensify all blue-light responses, but instead produces a specific pattern characterized by strong upregulation of metabolic and mitochondrial pathways together with suppression of interferon-associated signaling.

### OPN3 knockdown strongly attenuates the blue-light-dependent proteomic response

The contribution of OPN3 to the blue-light response was further assessed by comparing blue- and dark-exposed cells within the OPN3-silenced background. In the absence of atRAL, blue light induced only a minimal proteomic response in OPN3 knockdown cells, with 12 DEPs identified, all of which were upregulated (Fig. 7A–B; Table S1). Enrichment analysis showed that these proteins were mainly associated with vacuolar transport (NDFIP2, SQSTM1, NCOA4), intracellular iron ion homeostasis (NDFIP1, FTH1, NCOA4), positive regulation of autophagy (SQSTM1, CALCOCO2, TMEM59), and regulation of protein ubiquitination (NDFIP2, SQSTM1, NDFIP1) (Fig. 7B; Table S3).

**Fig. 7:**
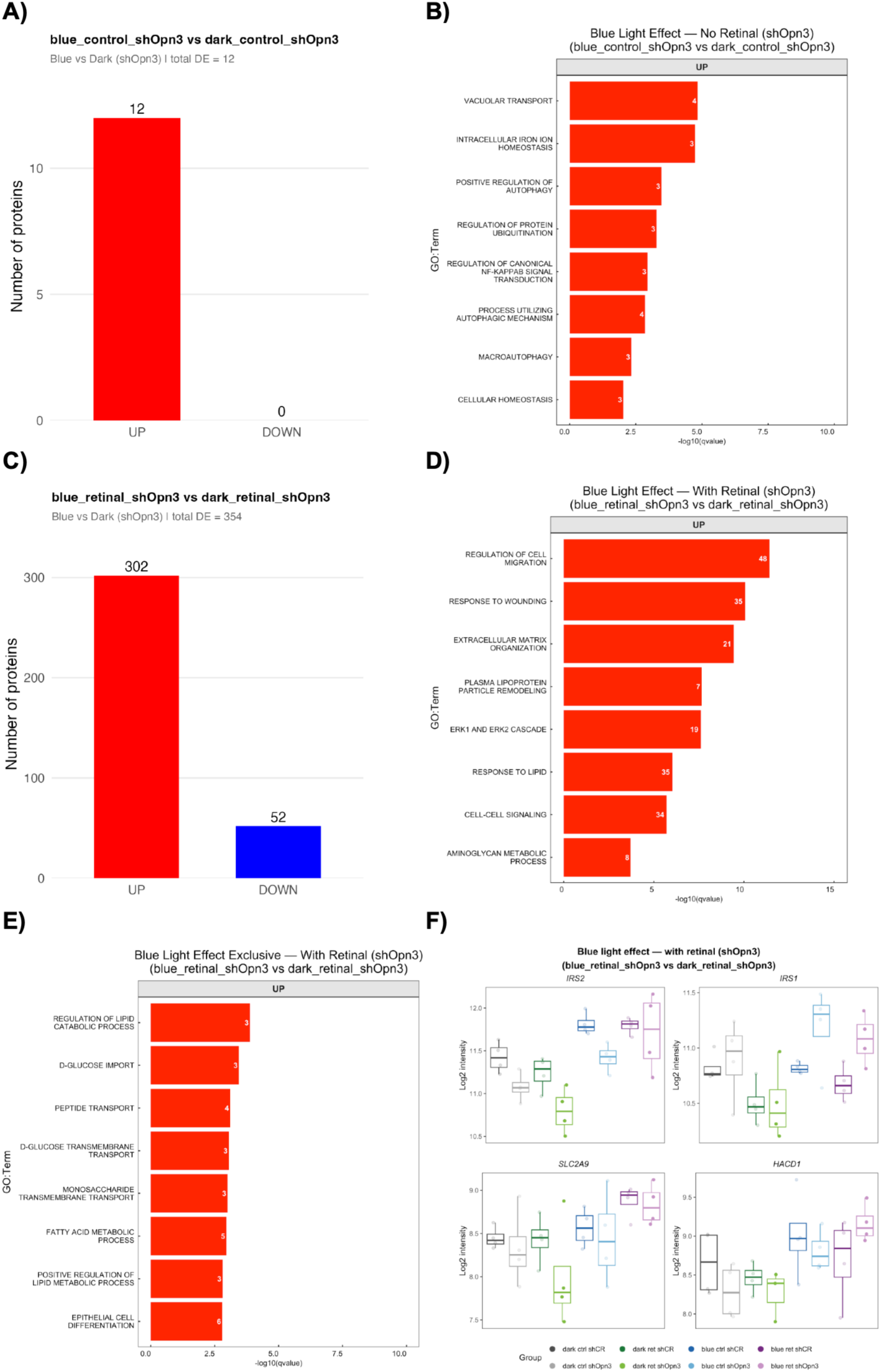
Blue light effects are dependent on OPN3 and its loss partially reduces retinal blue-light photosensitization. A and C) Histograms show the numbers of UP- and DOWN-regulated DEPs. B and D) Horizontal bar plots show the enriched biological processes according to identified DEPs. E) Exclusive biological processes identified in response to retinal photosensitization in OPN3 knockdown cells. F) Representative DEPs from the identified processes in E.

In the presence of atRAL, blue light still induced a broader response in OPN3-silenced cells, with 354 DEPs detected, including 302 upregulated and 52 downregulated proteins (Fig. 7C; Table S1). However, this response remained markedly lower than that observed in atRAL-supplemented shCR cells, where blue light induced 1547 DEPs. Enrichment of the upregulated proteins revealed biological processes related to blood vessel morphogenesis (HMOX1, MMP14, TGFB1), regulation of cell migration (IRS2, NRP1, EPHA2), response to wounding (SERPINE1, APOE, TGFB1), extracellular matrix organization (MMP14, TGFBI, FBLN1), ERK1/2 signaling (SPRY2, DUSP6, NRG1), and response to lipid (IRS1, PTGS2, PPARD) (Fig. 7D; Table S3).

Analysis of the exclusive DEPs further identified 44 proteins specific to the blue-light plus atRAL response in OPN3-silenced cells, including 33 upregulated and 11 downregulated proteins (Table S2). Enrichment of these exclusive proteins highlighted processes related to positive regulation of lipid catabolism (IRS2, IRS1, ABHD5), D-glucose import (IRS2, SLC2A9, IRS1), peptide transport (IRS2, SLC7A11, SLC25A39), organic anion transport (SLC7A11, SLC25A39, SLC2A9), and fatty acid metabolism (IRS2, ABHD5, HACD1) (Fig. 7E; Table S4). Representative proteins from this exclusive signature, including IRS1/IRS2, SLC2A9, SLC7A11, ABHD5, and HACD1, showed increased abundance specifically in the blue-light plus atRAL condition in OPN3-silenced cells (Fig. 7F).

Overall, these findings demonstrate that OPN3 is essential for the blue-light-dependent proteomic response in keratinocytes. In the absence of atRAL, OPN3 knockdown nearly abolishes the blue-light response, leaving only a small autophagy-, ubiquitination-, and iron-homeostasis-associated signature. In the presence of atRAL, blue light still triggers a residual response in OPN3-silenced cells, but this response is substantially reduced compared with shCR cells.

### OPN3 loss under blue light is associated with mitochondrial and metabolic suppression and limited additional atRAL-dependent remodeling

We next investigated the effect of OPN3 silencing under blue-light exposure, in the absence or presence of atRAL. In cells exposed to blue light without atRAL, OPN3 silencing induced a strong proteomic response, with 813 DEPs, including 137 upregulated and 676 downregulated proteins (Fig. 8A; Table S1). GO enrichment analysis showed that the downregulated DEPs were strongly associated with cellular respiration (SOD2, UQCRQ, ATP5PB), proton motive force-driven ATP synthesis (NDUFS2, ATP5PB, MT-ATP6), carbohydrate derivative biosynthetic process (POMK, B4GALT1, MGAT4B), macromolecule glycosylation (POMK, B4GALT1, MAN1A1), inflammatory response (S100A8, TNFAIP3, IL1B), lipid catabolic process (ACADM, CPT2, HADHA), and mitochondrial respiratory chain complex I assembly (NDUFS2, NDUFAF4, NDUFS8) (Fig. 8B; Table S3). Analysis of the exclusive DEPs further supported the idea that OPN3-silenced cells display a distinct blue-light-associated mitochondrial and metabolic vulnerability. In the blue-light condition without atRAL, 138 DEPs were exclusive to OPN3 silencing, including 38 upregulated and 100 downregulated proteins (Table S2). Exclusive GO enrichment was detected only among downregulated proteins and was mainly associated with mitochondrial gene expression (MTRF1L, GFM2, MRPS2), mitochondrial translation (GFM1, DARS2, MRPL27), mitochondrial RNA metabolic process (TRMT10C, FASTKD1, TWNK), amino acid activation/tRNA aminoacylation (YARS2, DARS2, AARS2), organic acid catabolic process (PCK2, CPT2, HADHA), cellular respiration/aerobic respiration (IDH3A, SUCLG1, COX6A1), energy derivation by oxidation of organic compounds (IDH3A, ETFA, MT-ATP6), and calcium import into the mitochondrion (HSPA9, AFG3L2, SLC25A23) (Fig. 8C–D; Table S4). These exclusive processes indicate that, under blue-light exposure, loss of OPN3 is associated with suppression of mitochondrial protein synthesis, respiratory metabolism, organic acid catabolism, and mitochondrial calcium-associated pathways.

**Fig. 8:**
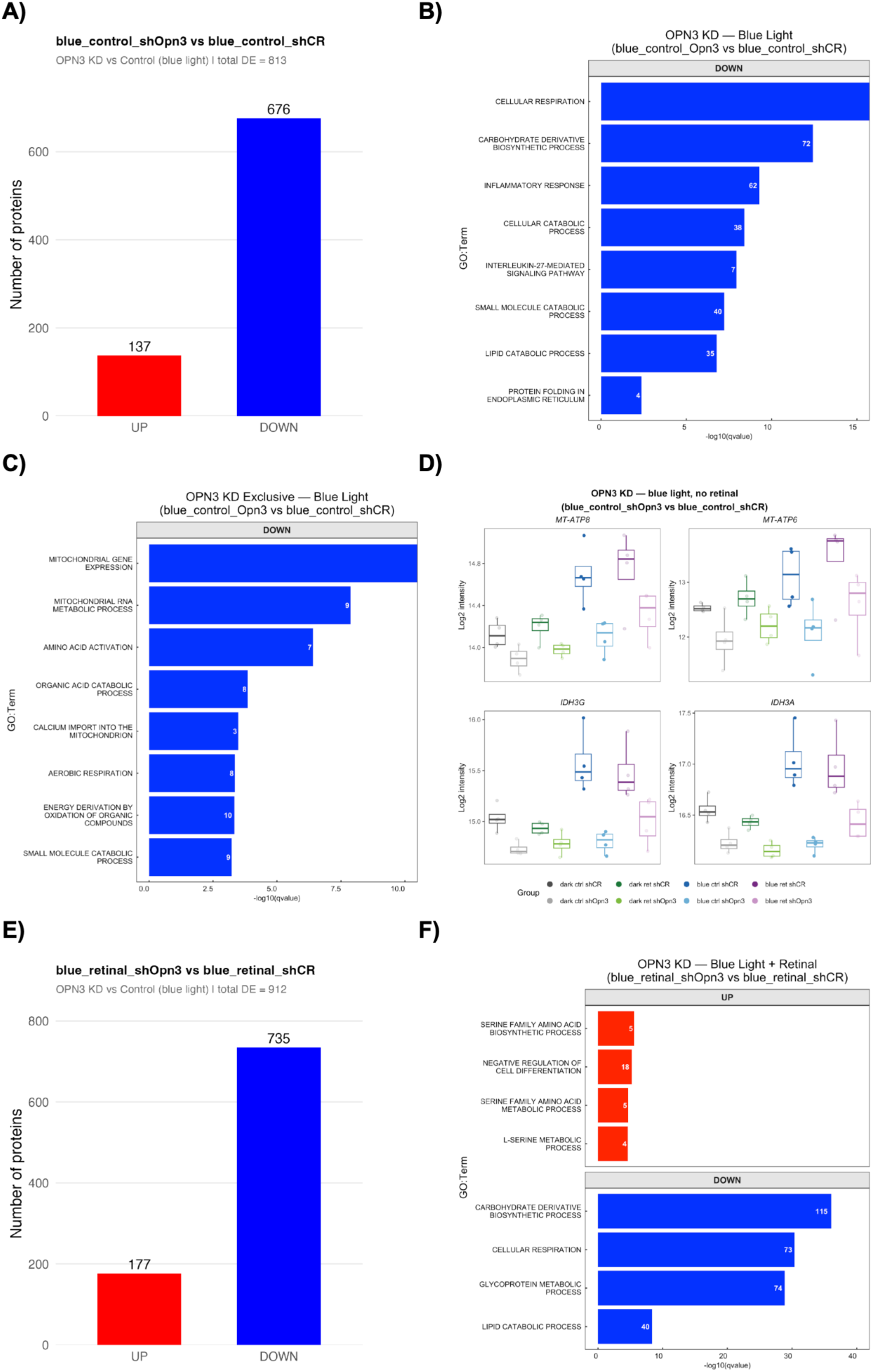
Loss of OPN3 makes cells prone to the blue light effect. A and E) Histograms show the numbers of UP- and DOWN-regulated DEPs. B – C) Horizontal bar plots show the enriched biological processes according to identified DEPs. D) Representative DEPs from the identified processes in C. F) Exclusive biological processes from DEPs identified in E.

When atRAL was present during blue-light exposure, OPN3 silencing induced a slightly larger proteomic response, with 912 DEPs, including 177 upregulated and 735 downregulated proteins (Fig. 8E; Table S1). Despite this increase, the overall biological signature remained highly similar and was again dominated by downregulation of mitochondrial, glycosylation, lipid-metabolic, and inflammatory pathways. Among upregulated DEPs, enriched processes were mainly related to serine family amino acid biosynthetic process (SHMT1, PSAT1, PSPH), serine family amino acid metabolic process (SHMT1, CTH, CBS), L-serine metabolic process (SHMT1, PSAT1, PSPH), and negative regulation of cell differentiation (LGALS1, RARG, GLI3) (Fig. 8F; Table S3). The downregulated DEPs in the blue-light plus atRAL condition were enriched in carbohydrate derivative biosynthetic process (MGAT4B, MOGS, TMEM165), glycoprotein metabolic process (MGAT4B, MOGS, B4GALT1), macromolecule glycosylation/glycosylation (MGAT4B, B4GALT1, POMK), cellular respiration (NDUFB6, COX4I1, ATP5PB), proton motive force-driven mitochondrial ATP synthesis (NDUFS2, ATP5PB, ATP5F1C), proton transmembrane transport (LETM1, ATP1B1, ATP5F1C), monoatomic ion homeostasis (SLC12A4, LETM1, ATP1B1), lipid catabolic process (ACADM, ACOX1, HSD17B4), lipid biosynthetic process (HSD17B2, ERG28, TSPO), and mitochondrial respiratory chain complex assembly (NDUFAF1, NDUFS2, SAMM50) (Fig. 8F; Table S3). Notably, the exclusive atRAL-associated effect in OPN3-silenced cells exposed to blue light was limited. Although 118 DEPs were exclusive to the blue-light plus atRAL OPN3-silencing comparison, including 58 upregulated and 60 downregulated proteins, exclusive GO enrichment identified only one biological process: serine family amino acid biosynthetic process (CTH, PSAT1, PSPH) (Table S2; Table S4). This suggests that atRAL adds a relatively narrow condition-specific component to an already strong OPN3-silencing signature under blue-light exposure.

Overall, these findings indicate that OPN3 loss under blue-light exposure is associated with broad suppression of mitochondrial respiration, mitochondrial protein synthesis, glycosylation, lipid metabolism, inflammatory signaling, and ion/proton homeostasis. The presence of atRAL modestly increases the number of DEPs but does not markedly modify the molecular signature.

### OPN3 downregulation increased recovery respiratory capacity after blue light exposure and atRAL photosensitization

Following proteomics predictions that suggested altered mitochondrial respiration, we measured the acute (0 h to 24 h) and recovery (48 h to 72 h) oxygen consumption rates (OCRs) in control and OPN3-knockdown cells after blue light and atRAL treatment. This experiment was designed to assess the acute effects of blue light stimulation in the presence or absence of atRAL, as well as the recovery (> 24 h) response of the cells. Blue light stimulation in the presence or absence of atRAL led to a marked suppression of respiration across the entire experiment (Fig. S7 A – B). In dark controls, no difference in OCR was observed between wild-type and OPN3-knockdown cells (Fig. S8A – C). In blue light-stimulated cells, the acute response in oxygen consumption was similar between groups in the presence or absence of atRAL (Fig. 9A – B). Interestingly, 72 h after blue light stimulation, OPN3-knockdown cells that were stimulated with atRAL and blue light showed increased oxygen consumption, indicating a faster recovery response (Fig. 9 A and C).

**Fig. 9:**
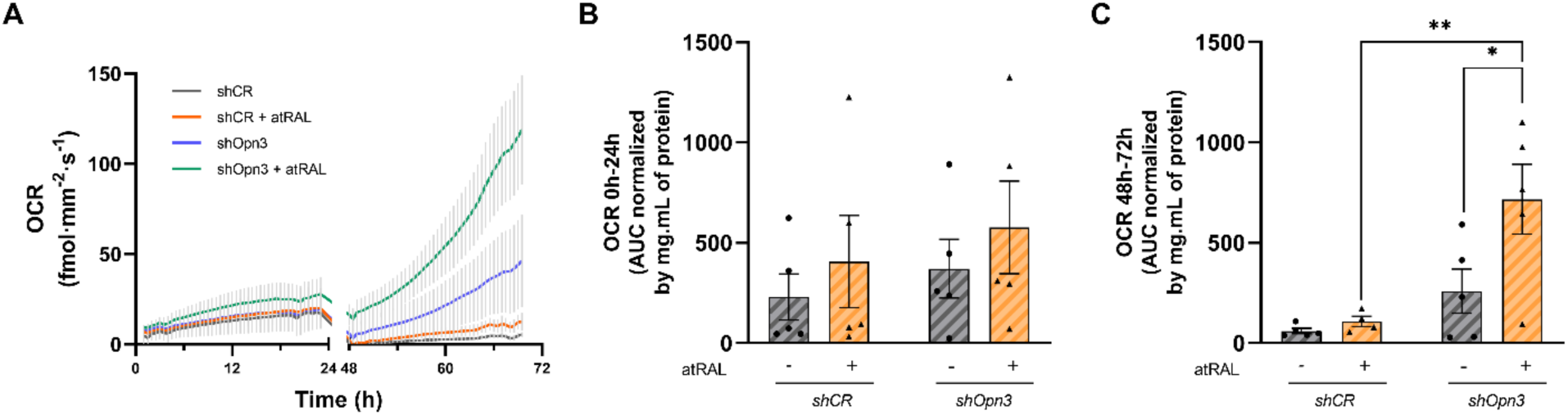
Oxygen consumption in OPN3-knockdown cells following atRAL photosensitization. (A) Oxygen consumption rates (OCRs) measured over 72 h in shCR and shOpn3 cells in the presence or absence of atRAL. (B) Quantification of OCR as the area under the curve (AUC) from 0–24 h, normalized by total protein content. (C) Quantification of OCR as the AUC from 48–72 h, normalized by total protein content. Each dot represents one independent biological replicate. Data are presented as the mean ± SEM from at least four independent experiments. Statistical analysis was performed using two-way ANOVA followed by Bonferroni’s multiple-comparisons test. ANOVA effects: OCR 0 h – 24 h (B)—interaction (*P* = 0.9390); OPN3-knockdown (*P* = 0.4173); retinal treatment (*P* = 0.3230). OCR 0 h – 72 h (C)—interaction (*P* = 0.0827); OPN3-knockdown (*P* = 0.0023); retinal treatment (*P* = 0.0373). \**P* < 0.05.

Overall, these findings show that the absence of OPN3 does not affect the blue-light response within the first twenty-four hours, but OPN3 knockdown cells show a high recovery respiratory response when stimulated with atRAL.

## DISCUSSION

Our findings establish OPN3 as a versatile regulator of cellular physiology by fulfilling three distinct yet interconnected roles. First, in the absence of atRAL or light exposure, OPN3 regulates pathways involved in autophagy, apoptosis, immune and interferon signaling, and cellular stress responses. Second, it is crucial for atRAL-dependent lipofuscin accumulation after blue-light exposure; silencing OPN3 prevents this accumulation without significantly affecting intracellular or extracellular atRAL levels. Third, OPN3 facilitates cellular adaptation to blue light independently of lipofuscin formation. Notably, atRAL dramatically enhances blue-light-induced proteomic remodeling in control cells, whereas OPN3 knockdown markedly diminishes this response. This attenuation is associated with widespread suppression of mitochondrial respiration, lipid metabolism, glycosylation, and ion-transport pathways. Collectively, these findings show that OPN3 supports basal cellular functions, enables atRAL-dependent lipofuscin formation, and sustains a protective metabolic program that shields keratinocytes from blue-light-induced stress.

### The effect of OPN3 as a light-independent sensor

Opsins were initially characterized as retinal photoreceptors and were later shown to mediate light-dependent responses in skin cells, including OPN3-dependent blue-light pigmentation ^22^, OPN4-dependent UVA and blue-light signaling ^37–40^, and OPN5-dependent UV-induced melanogenesis ^41^. Opsin functions are highly cell- and tissue-specific, extending from skin cells to multiple peripheral tissues ^42–44^. In *Drosophila larvae*, rhodopsins contribute to temperature discrimination and thermal preference ^45,46^. In mammalian spermatozoa, OPN2 and OPN4 activate distinct signaling pathways that guide thermotaxis ^47,48^, while in murine melanocytes and pre-adipocytes, OPN4 mediates temperature-dependent activation of circadian clock genes ^49,50^.

OPN3 has emerged as an important regulator of skin biology because it is expressed in major cutaneous cell types and has been functionally implicated in melanocytes, keratinocytes, and dermal fibroblasts ^21,51,52^. A major expansion of the OPN3 functional repertoire came in 2019, when Ozdeslik et al. showed that OPN3 negatively regulates melanogenesis independently of light through interaction with MC1R and inhibition of MC1R-dependent cAMP signaling ^53^. Subsequent studies identified basal OPN3 functions in melanocyte mitochondrial integrity and survival ^23^, TGFβ2-induced melanogenesis ^54^, BRAF^V600E^–ERK–MITF signaling ^55^, fibroblast iron and redox homeostasis and programmed cell death ^56^, epithelial stem-cell differentiation ^57^, and Langerhans-cell proliferation and migration ^58^. In hypothalamic neurons, constitutive OPN3 signaling suppresses MC4R-dependent cAMP production and potentiates Kir7.1 activity, thereby promoting food intake in mice ^59^. Consistent with these observations, silencing OPN3 in keratinocytes under dark, atRAL-free conditions led to an increase in proteins associated with apoptosis and chaperone-mediated autophagy, as well as several immune-related pathways, such as cytokine signaling and innate immune responses. In addition, Mao et al. (2025) demonstrated that OPN3 promotes inflammatory signaling following light exposure ^60^. Collectively, our findings further reveal that OPN3 maintains basal keratinocyte immune functions and proteome homeostasis independently of light and atRAL.

### The effect of OPN3 and atRAL on lipofuscin accumulation and the role of OPN3 as a light sensor

Our finding that blue-light photosensitization of atRAL induces lipofuscin accumulation represents a paradigm shift in understanding lipofuscin biology. To date, lipofuscin formation has been primarily attributed to: (i) accumulation of undegradable byproducts of oxidative damage, particularly in the retina where A2E and other bisretinoids are generated through the visual cycle ^16^; (ii) impaired autophagy leading to accumulation of damaged organelles and protein aggregates ^15^; and (iii) UVA-induced damage in keratinocytes ^9^. Here, we demonstrate that atRAL itself can serve as a photosensitizer that triggers lipofuscin accumulation when exposed to blue light. This is significant because atRAL is present in keratinocytes at physiologically relevant concentrations and can be generated from retinol, a widely used dermatological compound.

An increasing number of studies have implicated OPN3 in light-associated responses across several tissues. OPN3 was linked to blue-light-induced pigmentation in melanocytes ^22^, UVA-induced photoaging in dermal fibroblasts ^21^, light-dependent control of adipocyte metabolism and thermogenesis ^61,62^, and photorelaxation in vascular smooth muscle ^63,64^. More recently, OPN3 was reported to mediate blue-light-induced autophagy inhibition and pigmentation in melanocytes ^24^ and enhanced inflammatory signaling in keratinocytes in atopic dermatitis conditions ^60^. However, direct photochemical evidence remains largely limited to non-human OPN3 homologues ^65,66^. Most mammalian studies establish that OPN3 is required for a light response but do not demonstrate that it is the primary photon-absorbing receptor. Indeed, Ozdeslik et al. detected neither significant UV–visible absorption by human OPN3 nor an OPN3-dependent blue-light-induced Ca²⁺ response in melanocytes ^53^. Thus, OPN3 may function as a mediator or modulator of blue-light signaling, while its role as a direct photoreceptor remains unresolved. Consistent with this interpretation, OPN3 knockdown in our study markedly attenuated the blue-light proteomic response, reducing the number of DEPs from 284 to 12 in the absence of atRAL and from 1,547 to 354 in its presence. A particularly striking finding is that OPN3 knockdown reduces atRAL/blue-light-induced lipofuscin accumulation, despite not affecting atRAL uptake. This indicates that OPN3 plays an essential role in the pathway linking atRAL photosensitization to lipofuscin formation, but not simply as a transporter. The mechanism appears to involve a cascade: blue light photosensitizes atRAL, generating reactive species that impair mitochondrial function and lysosomal integrity, leading to autophagic flux inhibition. This is consistent with the mitochondrial-lysosomal axis theory of lipofuscin formation ^15^, but adds a novel initiating event: atRAL photosensitization. Importantly, this mechanism may be relevant beyond the skin, as atRAL is present in many tissues. Collectively, these findings identify OPN3 as a central mediator of blue-light responses in keratinocytes and show that its role in lipofuscin formation depends on atRAL photosensitization, without establishing OPN3 itself as the direct blue-light sensor.

### OPN3 supports a lipofuscin-independent adaptive response to blue light

Three studies provide precedent for an OPN3-dependent metabolic response to light. In brown adipocytes, Sato et al. showed that OPN3 supports both basal and light-stimulated energy metabolism: *Opn3*-deficient cells exhibited reduced glucose uptake and mitochondrial respiration even in darkness and failed to increase glucose utilization, CPT1 expression, and glucose- or fatty-acid-dependent respiration after light exposure ^62^. In white adipocytes, Nayak et al. demonstrated that blue light promotes OPN3-dependent lipolysis and hormone-sensitive lipase activation, thereby supplying fatty acids for thermogenesis; OPN3 loss reduced oxygen consumption, energy expenditure, and cold tolerance ^61^. More recently, Wu et al. showed that blue light stimulates lipid-droplet degradation in hepatocytes through OPN3-dependent PPARα activation and p62-mediated autophagic flux. OPN3 loss prevented PPARα nuclear accumulation, lipid clearance, ATP elevation, and the antiviral effects of blue light ^67^.

These studies closely parallel our proteomic findings and reveal an additional layer of complexity in the OPN3 response to blue light. Under blue-light exposure, OPN3 silencing was associated with broad suppression of cellular respiration, mitochondrial protein synthesis, ATP production, lipid metabolism, glycosylation, and ion/proton homeostasis. This signature was already evident without atRAL and remained largely similar after atRAL supplementation, indicating that it is predominantly independent of atRAL photosensitization and lipofuscin accumulation. At the same time, OPN3 knockdown markedly attenuated the overall blue-light proteomic response both without and with atRAL. Thus, OPN3 loss compromises mitochondrial and metabolic programs required for cellular adaptation to blue light response.

Our oxygen consumption assay showed that OPN3 knockdown does not affect OCR under dark conditions or during the 24 hours following irradiation. However, an interesting finding is that OPN3-knockdown cells recover faster 48-72 hours after irradiation, especially when treated with atRAL. We suggest these findings are associated with the overall low lipofuscin accumulation in OPN3-knockdown cells, but additional investigation is needed. Together, the literature and our findings support two partially separable functions of OPN3 under blue light: enabling atRAL-dependent lipofuscin accumulation and maintaining a lipofuscin-independent metabolic and autophagic adaptive program. The latter may involve constitutive signaling or protein–protein interactions, although its molecular basis remains to be determined.

## Conclusion

Our study has limitations: experiments were conducted in monolayer HaCaT keratinocyte cultures. Validation in primary keratinocytes and 3D skin models is required. Second, OPN3 was reduced by lentiviral knockdown, leaving the possibility of residual OPN3 activity. Third, the data evaluation pipeline applied predefined thresholds. Proteins falling slightly below either threshold may still contribute biologically. Consequently, the exclusive analyses highlight the most strongly affected proteins and pathways rather than completely condition-specific responses. Our blue light stimulation consists of a high dose (100 J/cm²). Lastly, although our findings identify OPN3 as a mediator of blue-light responses, they do not determine whether OPN3 directly senses blue light.

In conclusion, OPN3 contributes to keratinocyte homeostasis in at least three levels: it regulates basal immune, autophagic, and apoptosis-associated processes independently of light and atRAL, is required for atRAL-dependent lipofuscin accumulation under blue light and supports a largely atRAL-independent mitochondrial and metabolic adaptive program. These findings position OPN3 as a central regulator of both basal and blue-light-induced keratinocyte responses.

## MATERIALS AND METHODS

### Cell culture

Human immortalized non-malignant keratinocytes (HaCaT) were cultured in Dulbecco’s Eagle medium (DMEM) high-glucose, supplemented with 10% (v/v) fetal bovine serum (FBS), 1% streptomycin/penicillin (v/v), and 1 mM pyruvate and maintained in 37 °C incubator under a moist atmosphere of 5% carbon dioxide. Subculturing was performed twice a week, and the medium was replaced three times a week. Cells were routinely checked for mycoplasm contamination.

### Treatment with atRAL and irradiation protocol

For all experiments, cells were treated before irradiation with 5 µM of atRAL (Sigma-Aldrich) in culture medium for 30 minutes in a 37°C incubator under a moist atmosphere of 5% carbon dioxide. Control groups were incubated with the same volume of vehicle, resulting in a final ethanol concentration of 0.014% (v/v). After incubation, cells were washed two times with PBS and then irradiated with PBS. To avoid atRAL isomerization, all procedures were performed under low dim light without direct light exposure. Irradiation was performed with an LED-447 nm irradiator, provided with control of temperature set at 35°C (Ethik, Brazil). Before performing the irradiation, we measured the visible light irradiance (mW.cm^-2^) using a dosimeter (VLX-3W, France). To provide blue-light irradiation of 100 J.cm^-2^, HaCaT cells were irradiated for 77 minutes with 21 mW.cm^-2^ of irradiance. At the same time, the non-irradiated group was kept on a thermoblock at 35°C, protected from the light (dark group). After the irradiation protocol, PBS was replaced with culture medium, and cells were kept in culture conditions until the subsequent analysis. Treatment with atRAL and irradiation was always performed with cells at 50-60% confluence.

### Viability tests

For viability tests, 0.75×10^4^ cells per well were plated in 48-well plates and submitted to treatments when they reached 50-60% confluence. Mitochondrial viability, lysosomal viability, and membrane integrity were measured by MTT assay, neutral red assay, and staining with crystal violet, respectively. The cell viability for the three tests was calculated following the equation:

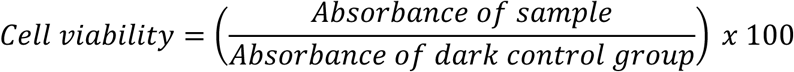

#### MTT

MTT was prepared at 5 mg.mL^-1^ in PBS, then cells were incubated with 0.05 mg. mL^-1^ of MTT diluted in culture medium for 2 hours. After this time, the medium was removed, and 200 µL of isopropanol was added to the wells. Plates were shaken at 300 rpm for 20 minutes at 25°C to dissolve crystals formed, and then absorbance was measured at 550 nm. Background was measured at 800 nm.

#### Neutral Red (NR)

Neutral red was prepared at 0.3 mg.mL^-1^ in PBS on the same day of use, then cells were incubated with 0.03 mg. mL^-1^ of neutral red diluted in culture medium for 2 hours. After this time, the medium was removed, cells were washed one time with PBS, and 100 µL of 1% acetic acid in 50% ethanol (v/v) was added. Plates were shaken at 300 rpm for 20 minutes at 25°C to dissolve crystals formed, and then absorbance was measured at 540 nm, and background was measured at 800 nm.

#### Crystal Violet Staining (CVS)

After neutral red measurements, wells were washed twice with PBS, and cells were stained with 0.02% of crystal violet for 5 minutes. After staining, cells were washed three times with PBS, and 200 µL of 0.1 M sodium citrate in 50% ethanol was added. Plates were shaken at 300 rpm for 30 minutes at 25°C to dissolve crystals formed, and then absorbance was measured at 585 nm.

### Lipofuscin quantification

#### Lipofuscin autofluorescence

Confocal microscopy was used to visualize lipofuscin granules. 1.5×10^4^ cells per well were plated in 24-well plates covered with glass for microscopy and submitted to treatments when they reached 50-60% confluence. 48 h after treatment, cells were washed two times with PBS, fixed with 4% paraformaldehyde for 15 minutes, and microscope slides were prepared with fluoromount-G with DAPI (Thermo Fisher Scientific). Images were acquired on a Leica confocal microscope (LAS X software, v3.5.7.23225) using an HC PL APO CS2 40×/1.30 oil-immersion objective. For detection of the lipofuscin autofluorescence signal, samples were excited with a 488 nm laser (White Light Laser line) and emission was collected between 509 and 601 nm using a hybrid detector (HyD), as previously described ^68^. DAPI was excited with a 405 nm diode laser and detected between 416 and 451 nm using a separate hybrid detector (HyD) channel, allowing identification of cell nuclei. The same laser power, detector settings, and acquisition parameters were maintained for all experimental groups. Fifty cells of each group were selected to measure integrated density relative to the area of the cell, expressing results as mean gray value.

### Autophagic flux integrity

#### Autophagy Arbitrary Units (AAU)

Autophagy was estimated using a mathematical formula that considers the relationship between lysosomal viability, mitochondrial viability, and membrane integrity, and is expressed as autophagy arbitrary units (AAU) ^32^. To calculate AAU, the mean NR cell viability was normalized to the mean of the MTT and CVS cell viability rates according to the function w (x, y, z):

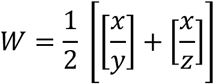

Where x, y, and z are the survival rates measured by NR, CVS, and MTT assays, respectively.

#### LC3AB-II net flux

Immunocontent of LC3-II was measured 24 hours after treatments by Western Blot. In the LC3-II net-flux degradation of LC3-II inside the autolysosome is estimated by the comparison of two samples with and without lysosomal inhibitor treatment. In this case, for each condition, there was another paired group where 200 nM of bafilomycin was added 1 h before protein extraction. The differences in the amount of LC3-II between samples in the presence and absence of bafilomycin represent the amount of LC3-II that is delivered to lysosomes for degradation (i.e., autophagic flux) ^33,34^.

### Western Blot

For Western Blot analysis, 0.6×10^6^ cell per well were plated in 6-well plates and submitted to treatments when reached 50-60% confluence. 24 hours after treatment, cells were washed twice with cold PBS and lysed with 100 µL of RIPA lyses buffer (Thermo Fisher Scientific) with 1% of protease inhibitor and phosphatase cocktail (Sigma-Aldrich). Plates were kept on ice for 5 minutes; after that, cell scrapers were used to remove all cells from the well. Samples were centrifuged at 14000 *g* for 15 minutes at 4°C and supernatant was used for analysis. Protein samples were mixed with 1% mercaptoethanol and sample loading buffer and denatured for 5 minutes at 95°C. 30 µg of total proteins were separated by 15 % sodium dodecyl sulfate–polyacrylamide gel electrophoresis (SDS–PAGE) and transferred on to polyvinylidene difluoride (PVDF) membranes (Amersham® Hybond P 0.20) overnight at 25 v. Membranes were blocked in Tris-buffered saline (TBS; 50 mM Tris-HCl, pH 7.6, 150 mM NaCl) with 5% (w:v) non-fat milk and 0.1% (v:v) Tween® 20 for 1h at room temperature and incubated with primary monoclonal antibodies in TBS with 2.5% (w:v) BSA and 0.1% (v:v) Tween® 20 overnight at 4°C. For WB, rabbit anti-human LC3A/B monoclonal antibody (Cell signaling, 12741; 1:500) and rabbit anti-human LAMP-1 monoclonal antibody (Cell signaling, 9091; 1:1000) were used as the primary antibody. Mouse anti-human beta actin monoclonal antibody (abcam, ab6276; 1:10000) was used as housekeeping. Three washing steps in TBS with 0.1% (v:v) Tween® 20 for 10 min were performed and followed by incubation with fluorescent anti-rabbit IRDye 680RD (Li-cor 926-68071; 1:5000) or anti-mouse IRDye 800CW (Li-cor 926-32210; 1:20000) secondary antibody diluted in TBS with 2.5% (v:v) BSA and 0.1% (v:v) Tween® 20 for 1h at room temperature. Membranes were washed twice in TBS with 0.1% (v:v) Tween® 20 and once in TBS, 10 min each. Fluorescence was detected using the Odyssey CLx Imager (Li-cor) and band densitometry was quantified using Fiji ImageJ software.

### Labeling of lysosomes and acidic vacuoles in live cells

0.1×10^6^ cells per well were plated in 35 mm dishes with four compartments with bottom glass and submitted to treatments when they reached 50-60% confluence. Three and 24 h after treatment, cells were incubated for 30 minutes with Lysotracker deep red (50 nM of) (Thermo Fisher Scientific) or acridine orange (1 µg/µL) (Acros Organics) in culture medium. After this time, the culture medium was replaced by PBS, and immunofluorescence microscopy was performed with live cells. For lysotracker deep red, samples were excited at 590-650 nm and emission was detected at 662-738 nm (Y5). 50 cells of each group were selected to measure the integrated density relative to the area of the cell. Results are expressed as fold change relative to the dark control group. For acridine orange, fluorescence was detected in two channels. Samples were excited at 460-500 nm, and emission was detected at 512-542 nm (FITC) or excited at 541-551 nm and emission was detected at 565-605 nm (RHOD). The ratio of acridine orange emission at RHOD to emission at FITC was used to quantify the increase in the number of acidic vesicular organelles observed during cellular autophagy. 50 cells of each group were selected to measure the integrated density relative to the area of the cell. Results are expressed as fold change relative to the dark control group.

### Gene expression – RT q-PCR

0.6×10^6^ cells per well were plated in 6-well plates and submitted to treatments when reached 50-60% confluence. Six hours after treatment, RNA was extracted and purified with PureLink RNA Mini Kit (Thermofisher). Integrity was checked by capillary electrophoresis (Bioanalyzer). 2 µg of RNA was used to produce cDNA (High-Capacity cDNA Reverse Transcription Kit, Thermofisher). Only samples with A260/280 higher than 1.8 and RIN higher than 7 were used. RT-qPCR was performed following SYBR Green PCR Master Mix instructions (Life Technologies 4309155). For OPN3 expression, primers were F-CCTGGTCAACATCAGCCTCA and R-GGCAATGGAAACAATCCCGAAG. Beta-actin was used as housekeeping, primers F-CACAGAGCCTCGCCTTTGC and R-AATCCTTCTGACCCATGCCC. ΔCt was normalized by housekeeping and results are expressed as ΔΔCt.

### Silencing protocol

To produce knockdown cells for OPN3, 0.2×10^6^ cells per well were plated in 12-well plates. On the next day, cells were incubated overnight with 8 µg/mL of Polybrene and 15 µL of viral particles (sc-45989-V or sc-108080 – Santa Cruz) for transduction of cells. On the third day, polybrene was removed and the medium culture was replaced. Transduced cells were selected with 2 µg/mL of puromycin and the puromycin-selected cell population, rather than individual clones, was used in all subsequent experiments. The efficiency of transduction was confirmed by RT-qPCR.

### atRAL extraction and quantification

Retinoids were extracted following a previously proposed method ^69^. shCR and shOpn3 cells were seeded at 3×10^6^ in T75 flasks and submitted to treatment with atRAL or vehicle when reached 70-80% of confluence. Immediately after atRAL incubation, cells were washed three times with PBS, detached with trypsin EDTA 0.25%, centrifuged at 1500 g for 10 minutes, and resuspended in 200 µL of PBS. Then, 200 µL of ethanol, 90 µL of 2 M sodium hydroxide, and 245 µL of n-hexane were added to the samples. Samples were shaken at 200 rpm for 10 minutes at 30°C, centrifuged at 3000 g for 5 minutes, and the upper layer fraction containing retinoids was collected. The extraction with n-hexane was repeated two more times to increase the extraction efficiency. After this, n-hexane evaporated with nitrogen gas, and samples were resuspended in 300 µL of ethanol. The atRAL absorption was measured at 383 nm. Non-treated samples were used as the blank for their respective groups. Results are expressed as fold change in relation to shCR group. All procedures were performed in the dark without direct light contact to avoid retinal isomerization.

### Proteomics sample processing and data processing

Proteins were extracted in lysis buffer (2% sodium dodecyl sulfate, 50mM HEPES) using focused ultra-sonication (Covaris ML230) followed by addition of approximately 0.5% Benzonase, vortex and shaking until the cell pellet was no longer visible. Protein concentrations were determined using Pierce BCA Protein Assay Kit (Thermo Scientific) on a SpectraMax iD3 microplate reader (Molecular Devices). Samples were processed with a modified SP3 method ^70^ using a Beckman i7 robotic system equipped with a magnet, shaker and incubator. Samples (100 µg protein) were reduced in 10mM dithiothreitol at 60°C for 30 min during shaking and alkylated with 20mM chloroiodoacetamide at room temperature for 10min during shaking. Washed hydrophobic and hydrophilic Sera-Mag™ SpeedBeads (Carboxylate-Modified, Cytiva) were added to the samples with a bead to protein ratio of 7:1. Proteins were precipitated on the beads by acetonitrile (ACN) (final concentration 70%), washed two times with 70% ethanol, one wash with ACN and dried at room temperature. Beads were resuspended in 140 µL 50 mM HEPES, 10mM CaCl2,10% ACN and proteins were digested with Rapizyme trypsin [1:40] (Waters Corporation) for two hours and for a second time [1:40] overnight incubation at 37°C. The next day, milli-Q water was added, mixed, and beads were shaked. Peptides were cleaned using precipitation on the beads with a final concentration of 95% ACN during shaking. Beads were washed three times with 100% ACN. Peptides were extracted with milliQ water and transferred to a clean 96-well plate. An extra clean-up step were performed, after addition of trifluoroacetic acid (TFA) to lower pH below 2, using a trace-N20 NBE 5mg (Tecan) 96-well plate, washed with ACN and 0.1% TFA, loading the samples three times, washed with 0.1% TFA and eluted twice with 50% ACN. Peptides were dried using vacuum centrifugation and redissolved in 3% ACN, 0.1% TFA, 0.015% DDM (n-dodecyl -D-maltoside). Peptide concentrations were determined using Pierce™ Quantitative Fluorometric Peptide Assay (Thermo Scientific). Samples were normalized to 0.015 ug/ul in 3% ACN, 0.1% TFA, 0.015% DDM and 300 ng peptides were loaded on Evotips according to the manufacturer’s instructions.

### Label-free Data-Independent Acquisition (DIA) quantification LC-MS/MS analysis

The samples were analysed on an Orbitrap Astral mass spectrometer (Thermo Fisher Scientific) coupled to an EvoSep Eno LC system (EvoSep Biosystems). Peptides (300 ng) were separated on a PepSep column (8 cm x 150 μm) using a 60 samples per day (60 SPD) method. The precursor ion mass spectra were acquired at a resolution of 240 000, an m/z range of 380-980 and a normalized AGC target of 500%. Astral DIA standard window settings were used with an isolation window of 2 m/z, 3.5 ms scan time, 299 scan events, cycle time of 0.6 s, normalized higher-energy collisional dissociation (HCD) energy of 27 and a customized AGC target of 500%.

Data analysis was performed with Spectronaut (20.5, Biognosys) using directDIA workflow. The data was matched against the reviewed human database (downloaded from uniprot.org, 20421 entries, January 2025, including pig trypsin). Standard settings were used, where the maximum missed cleavage rate was set to 1 and Proteotypicity Filter set to Only Protein Group Specific, run-level protein scoring was set to Highest Scoring Observation and stringent identification cutoff of 0.01 (precursor q-value cutoff (experiment, run), precursor posterior error probability (PEP) cutoff, protein q-value cutoff (experiment/run) and protein PEP cutoff) settings were used as described by ^71^.

### Proteomics bioinformatic workflow

#### Data normalization

Protein group intensity data were exported from the DIA analysis software as software-normalized MS2-level and imported into R for downstream preprocessing. Proteins were then filtered for quantitative completeness within each experimental group: a protein was retained only if it presented no more than one missing value out of four biological replicates in every group, corresponding to a maximum missingness of 25% per group. A total of 9,178 unique proteins were retained after filtering. Retained intensity values were log₂-transformed and normality was confirmed using Q-Q plots. Homogeneity of variance across groups was assessed per protein using Levene’s test, with p-values BH-adjusted for multiple testing. 0.1% of the proteins showed significant heteroscedasticity, supporting standard limma empirical Bayes shrinkage without group-specific variance modeling.

#### Differentially expressed proteins

Differentially abundant proteins were identified using the limma ^72^ package in R. A linear model was fitted using a cell-means parameterization with no intercept (∼ 0 + group), in which each coefficient represents the mean log2 intensity of one experimental group. The base model was fitted with lmFit(), and empirical Bayes variance moderation was applied after contrast fitting using eBayes() to stabilize protein-wise variance estimates. As an overview of global proteome variation, a moderated F-test was performed by defining seven contrasts comparing each non-reference group against the common reference group (dark_control_shCR). These contrasts were fitted from the base model using contrasts.fit(), followed by empirical Bayes moderation with eBayes(), and a multi-contrast F-statistic was extracted using topTable(). This global F-test was used as a descriptive summary of proteins varying across experimental groups.

Primary differential abundance analysis was performed using pre-specified pairwise contrasts within the 2 × 2 × 2 factorial design. The pairwise contrasts were fitted from the base model using a single contrast matrix, followed by empirical Bayes moderation with eBayes(). For each contrast, p-values were adjusted across proteins using the Benjamini–Hochberg procedure. A protein was considered differentially expressed if it met both an FDR threshold of < 0.05 and an absolute log2 fold change > 0.58. To identify proteins with contrast-specific responses, an exclusivity analysis was performed for each of the twelve pairwise contrasts. A protein was classified as exclusive to a given contrast if it was not detected as differentially abundant in any of the remaining seven pairwise contrasts.

### Enrichment analyses

Gene Ontology (GO) Biological Process enrichment was performed for each pairwise contrast using the clusterProfiler package (enrichGO()), with org.Hs.eg.db as the annotation source and gene symbols (keyType = “SYMBOL”) used directly. The background universe was defined as all identified proteins. Enrichment was tested separately for up- and down-regulated proteins within each contrast. Resulting terms were further filtered to require a q-value < 0.05 and a minimum protein count of 3 per term. For contrasts yielding more than 100 significant terms, redundant GO terms were collapsed using clusterProfiler::simplify() (cutoff = 0.6). This enrichment procedure was applied both to the full DEP sets per contrast and, separately, to the subset of DEPs classified as exclusive to each contrast.

### Long-term Oxygen Consumption Rates (OCR)

To measure the effects of blue light irradiation and atRAL treatment in OPN3-knockdown cells oxygen consumption rates (OCRs) were measured in cells cultured under standard culture conditions using a Resipher Real-time Cell Analyzer (Lucid Scientific, Atlanta, GA, USA) for three consecutive days. Briefly, 3 × 10⁴ cells per well were seeded in a 96-well plate and allowed to adhere for 24 h under standard culture conditions. Following atRAL treatment and blue light irradiation, as described above, the PBS was replaced with phenol red-free DMEM supplemented with 25 mM glucose and 10% FBS, and the Resipher sensor system was immediately positioned over the plate. Basal OCRs were monitored continuously for three consecutive days, with three technical replicates per experimental condition. Cell-free wells containing the same culture medium were included to correct for background. The culture medium was replaced daily. At the end of each measurement day, total protein content was quantified, and the values were used to normalize the area under the curve (AUC) of the OCR measurements.

## DATA AVAILABILITY STATEMENT

The datasets generated and/or analyzed during the current study are available from the corresponding author upon request. The mass spectrometry proteomics data have been deposited to the ProteomeXchange Consortium ^73^ via the MassIVE ^74^ repository with the dataset identifiers PXD083694 and MSV000103113.

## ACKNOWLEDGEMENTS

We would like to thank the Fundação de Amparo à Pesquisa do Estado de São Paulo (FAPESP) grants (CEPID -Redoxoma, 2013/07937-8, 2024/17859-9) for funding the projects. de Assis, LVM received support from the Knut and Alice Wallenberg Foundation as a Wallenberg Molecular Medicine Fellow, the Jeanssons Foundation, and the German Research Foundation (DFG) under grant TRR 418 (ID 541063275, B04). The Proteomics Core Facility, Sahlgrenska Academy, University of Gothenburg has financial support from SciLifeLab and BioMS national infrastructures (Swedish Research Council, VR). The authors thank Stina Lassessonand Elham Rekabdarfor their technical support.

## AUTHOR CONTRIBUTIONS STATEMENT (CREDIT-COMPLIANT)

Conceptualization: MVD, LVMA, MSB

Methodology: MVD, HCJ, MAHL, JF, KT, CSW

Validation: MVD, LVMA, MSB

Formal analysis: MVD, GL, LVMA, MSB

Investigation: MVD, MAHL

Resources: MSB and LVMA

Data Curation: MVD, GL, LVMA

Writing - Original Draft: MVD and LVMA

Writing - Review & Editing: All authors

Supervision: LVMA and MSB

Project administration: MVD, LVMA, MSB

Funding acquisition: CSW, LVMA, and MSB

## ARTIFICIAL INTELLIGENCE STATEMENT

The bioinformatic pipeline was developed by the authors and improved using ChatGPT and Claude. Grammarly was used to improve the clarity of the manuscript. The authors take full responsibility for the accuracy, integrity, and reproducibility of the manuscript.

## REFERENCES

1. Liebel, F., Kaur, S., Ruvolo, E., Kollias, N. & Southall, M. D. Irradiation of skin with visible light induces reactive oxygen species and matrix-degrading enzymes. J. Invest. Dermatol. 132, 1901–1907 (2012).

2. Nakashima, Y., Ohta, S. & Wolf, A. M. Blue light-induced oxidative stress in live skin. Free Radic. Biol. Med. 108, 300–310 (2017).

3. Tonolli, P. N., Marie, S. K. N., Oba-Shinjo, S. M., de Assis, L. V. M. & Baptista, M. S. Stage-specific phenotypic and transcriptional alterations in HaCaT keratinocytes exposed to acute and chronic blue light. Photochem. Photobiol. 101, 947–959 (2025).

4. de Assis, L. V. M., Tonolli, P. N., Moraes, M. N., Baptista, M. S. & de Lauro Castrucci, A. M. How does the skin sense sun light? An integrative view of light sensing molecules. J. Photochem. Photobiol. C Photochem. Rev. 47, 100403 (2021).

5. Austin, E. et al. Visible light. Part I: Properties and cutaneous effects of visible light. J. Am. Acad. Dermatol. 84, 1219–1231 (2021).

6. Jeong, S. Y., Gu, X. H. & Jeong, K. W. Photoactivation of N-retinylidene-N-retinylethanolamine compromises autophagy in retinal pigmented epithelial cells. Food Chem. Toxicol. 131, (2019).

7. Santacruz-Perez, C., Tonolli, P. N., Ravagnani, F. G. & Baptista, M. S. Photochemistry of Lipofuscin and the Interplay of UVA and Visible Light in Skin Photosensitivity. in Photochemistry and Photophysics - Fundamentals to Applications (InTech, 2018). doi:10.5772/intechopen.76641.

8. Tonolli, P. N. et al. Lipofuscin in keratinocytes: Production, properties, and consequences of the photosensitization with visible light. Free Radic. Biol. Med. 160, 277–292 (2020).

9. Tonolli, P. N. et al. Lipofuscin Generated by UVA Turns Keratinocytes Photosensitive to Visible Light. J. Invest. Dermatol. 137, 2447–2450 (2017).

10. Anstötz, M. et al. Palmitoyl-Protein Thioesterase 1 (PPT1) Protein, Linked to Neuronal Ceroid Lipofuscinosis 1, Is a Major Constituent of Ageing-Related Human Neuronal Lipofuscin. Neuropathol. Appl. Neurobiol. 51, (2025).

11. Baldensperger, T. et al. The age pigment lipofuscin causes oxidative stress, lysosomal dysfunction, and pyroptotic cell death. Free Radic. Biol. Med. 225, 871–880 (2024).

12. Tieze, S. M. et al. Molecular elucidation of brain lipofuscin in aging and Neuronal Ceroid Lipofuscinosis. Res. Sq. https://doi.org/10.21203/RS.3.RS-6010379/V1 (2025) doi:10.21203/RS.3.RS-6010379/V1.

13. Mitter, S. K. et al. Autophagy in the Retina: A Potential Role in Age-Related Macular Degeneration. Adv. Exp. Med. Biol. 723, 83 (2012).

14. Shin, Y. C. et al. A small molecule compound that inhibits blue light-induced retinal damage via activation of autophagy. Biochem. Pharmacol. 211, (2023).

15. Brunk, U. T. & Terman, A. The mitochondrial-lysosomal axis theory of aging: Accumulation of damaged mitochondria as a result of imperfect autophagocytosis. Eur. J. Biochem. 269, 1996–2002 (2002).

16. Sparrow, J. R. et al. A2E, a byproduct of the visual cycle. Vision Res. 43, 2983–2990 (2003).

17. Crouch, R. K., Koutalos, Y., Kono, M., Schey, K. & Ablonczy, Z. A2E and Lipofuscin. Prog. Mol. Biol. Transl. Sci. 134, 449–463 (2015).

18. VanBuren, C. A. & Everts, H. B. Vitamin A in Skin and Hair: An Update. Nutrients 14, (2022).

19. Choi, E. H., Daruwalla, A., Suh, S., Leinonen, H. & Palczewski, K. Retinoids in the visual cycle: role of the retinal G protein-coupled receptor. J. Lipid Res. 62, (2021).

20. Castellano-Pellicena, I. et al. Does blue light restore human epidermal barrier function via activation of Opsin during cutaneous wound healing? Lasers Surg. Med. 51, 370–382 (2019).

21. Lan, Y., Wang, Y. & Lu, H. Opsin 3 is a key regulator of ultraviolet A-induced photoageing in human dermal fibroblast cells. Br. J. Dermatol. 182, 1228–1244 (2020).

22. Regazzetti, C. et al. Melanocytes Sense Blue Light and Regulate Pigmentation through Opsin-3. J. Invest. Dermatol. 138, 171–178 (2018).

23. Wang, Y., Lan, Y. & Lu, H. Opsin3 Downregulation Induces Apoptosis of Human Epidermal Melanocytes via Mitochondrial Pathway. Photochem. Photobiol. 96, 83–93 (2020).

24. Yu, E. et al. The Pigmentation of Blue Light Is Mediated by Both Melanogenesis Activation and Autophagy Inhibition through OPN3–TRPV1. J. Invest. Dermatol. 145, 908–918.e6 (2025).

25. Adler, L. et al. The 11-cis Retinal Origins of Lipofuscin in the Retina. Prog. Mol. Biol. Transl. Sci. 134, e1–e12 (2015).

26. Balado-Simó, P. et al. An Updated Review of Topical Tretinoin in Dermatology: From Acne and Photoaging to Skin Cancer. J. Clin. Med. 14, (2025).

27. Dragicevic, N. & Maibach, H. I. Liposomes and Other Nanocarriers for the Treatment of Acne Vulgaris: Improved Therapeutic Efficacy and Skin Tolerability. Pharmaceutics 16, (2024).

28. Maalmi, H. et al. Dose-Response Relationship between Serum Retinol Levels and Survival in Patients with Colorectal Cancer: Results from the DACHS Study. Nutrients 10, (2018).

29. Yilmaz, G. et al. Baseline serum vitamin A and vitamin C levels and their association with disease severity in COVID-19 patients. Acta Biomed. 94, (2023).

30. Chihara, K. & Waddell, W. H. Electronic and Vibrational Spectral Investigation of the Molecular Association of the All-Trans Isomers of Retinal, Retinol, and Retinoic Acid. J. Am. Chem. Soc. 102, 2963–2968 (1980).

31. Horwitz, J. & Heller, J. Interactions of all trans, 9, 11, and 13 cis retinal, all transretinyl acetate, and retinoic acid with human retinol binding protein and prealbumin. J. Biol. Chem. 248, 6317–6324 (1973).

32. Martins, W. K. et al. Parallel damage in mitochondria and lysosomes is an efficient way to photoinduce cell death. Autophagy 15, 259–279 (2019).

33. Mizushima, N., Yoshimori, T. & Levine, B. Methods in Mammalian Autophagy Research. Cell 140, 313–326 (2010).

34. Yamamoto-Imoto, H., Hara, E., Nakamura, S. & Yoshimori, T. Measurement of autophagy via LC3 western blotting following DNA-damage-induced senescence. STAR Protoc. 3, 101539 (2022).

35. Cheng, X. T. et al. Revisiting LAMP1 as a marker for degradative autophagy-lysosomal organelles in the nervous system. Autophagy 14, 1472–1474 (2018).

36. Eskelinen, E. L. Roles of LAMP-1 and LAMP-2 in lysosome biogenesis and autophagy. Mol. Aspects Med. 27, 495–502 (2006).

37. Kusumoto, J. et al. OPN4 belongs to the photosensitive system of the human skin. Genes to Cells 25, 215–225 (2020).

38. de Assis, L. V. M., Moraes, M. N., Magalhães-Marques, K. K. & Castrucci, A. M. de L. Melanopsin and rhodopsin mediate UVA-induced immediate pigment darkening: Unravelling the photosensitive system of the skin. Eur. J. Cell Biol. 97, 150–162 (2018).

39. de Assis, L. V. M., Moraes, M. N., da Silveira Cruz-Machado, S. & Castrucci, A. M. L. The effect of white light on normal and malignant murine melanocytes: A link between opsins, clock genes, and melanogenesis. Biochim. Biophys. Acta - Mol. Cell Res. 1863, 1119–1133 (2016).

40. de Assis, L. V. M. et al. Melanopsin mediates UVA-dependent modulation of proliferation, pigmentation, apoptosis, and molecular clock in normal and malignant melanocytes. Biochim. Biophys. Acta - Mol. Cell Res. 1867, 118789 (2020).

41. Lan, Y., Zeng, W., Dong, X. & Lu, H. Opsin 5 is a key regulator of ultraviolet radiation-induced melanogenesis in human epidermal melanocytes. Br. J. Dermatol. 185, 391 (2021).

42. Castrucci, A. M. de L., Baptista, M. S. & de Assis, L. V. M. Opsins as main regulators of skin biology. *J*. Photochem. Photobiol. 15, 100186 (2023).

43. Moraes, M. N., de Assis, L. V. M., Provencio, I. & Castrucci, A. M. de L. Opsins outside the eye and the skin: a more complex scenario than originally thought for a classical light sensor. Cell Tissue Res. 385, 519–538 (2021).

44. Andrabi, M., Upton, B. A., Lang, R. A. & Vemaraju, S. An Expanding Role for Nonvisual Opsins in Extraocular Light Sensing Physiology. Annu. Rev. Vis. Sci. 9, 245–267 (2023).

45. Shen, W. L. et al. Function of rhodopsin in temperature discrimination in Drosophila. Science 331, 1333–1336 (2011).

46. Sokabe, T., Chen, H. C., Luo, J. & Montell, C. A Switch in Thermal Preference in Drosophila Larvae Depends on Multiple Rhodopsins. Cell Rep. 17, 336–344 (2016).

47. Pérez-Cerezales, S. et al. Involvement of opsins in mammalian sperm thermotaxis. Sci. Rep. 5, (2015).

48. Roy, D., Levi, K., Kiss, V., Nevo, R. & Eisenbach, M. Rhodopsin and melanopsin coexist in mammalian sperm cells and activate different signaling pathways for thermotaxis. Sci. Rep. 10, 112 (2020).

49. Moraes, M. N. et al. Melanopsin, a Canonical Light Receptor, Mediates Thermal Activation of Clock Genes. Sci. Reports 2017 71 7, 13977-(2017).

50. Zanetti, G., de Assis, L. V. M., Moraes, M. N. & de Lauro Castrucci, A. M. Melanopsin mediates thermal activation of molecular clock genes in murine preadipocytes. Chronobiol. Int. https://doi.org/10.1080/07420528.2026.2679133 (2026) doi:10.1080/07420528.2026.2679133.

51. Haltaufderhyde, K., Ozdeslik, R. N., Wicks, N. L., Najera, J. A. & Oancea, E. Opsin Expression in Human Epidermal Skin. Photochem. Photobiol. 91, 117–123 (2015).

52. Olinski, L. E., Lin, E. M. & Oancea, E. Illuminating insights into opsin 3 function in the skin. Adv. Biol. Regul. 75, (2020).

53. Ozdeslik, R. N., Olinski, L. E., Trieu, M. M., Oprian, D. D. & Oancea, E. Human nonvisual opsin 3 regulates pigmentation of epidermal melanocytes through functional interaction with melanocortin 1 receptor. Proc. Natl. Acad. Sci. U. S. A. 116, 11508–11517 (2019).

54. Wang, Y., Lan, Y., Yang, X., Gu, Y. & Lu, H. TGFβ2 Upregulates Tyrosinase Activity through Opsin-3 in Human Skin Melanocytes In Vitro. J. Invest. Dermatol. 141, 2679–2689 (2021).

55. Dong, X. et al. OPN3 Regulates Melanogenesis in Human Congenital Melanocytic Nevus Cells through Functional Interaction with BRAFV600E. J. Invest. Dermatol. 142, 3020–3029.e5 (2022).

56. Liu, T. et al. Downregulation of OPN3 Gene Induces Ferroptosis, Apoptosis, and Pyroptosis of Human Dermal Fibroblasts In Vitro. J. Invest. Dermatol. 144, 2305–2308.e6 (2024).

57. Jin, S. et al. In vitro differentiation of human amniotic epithelial stem cells into keratinocytes regulated by OPN3. Exp. Dermatol. 33, (2024).

58. Ye, T. et al. Opsin 3 expression in human Langerhans cell histiocytosis and its mediation of ELD-1 cellular function. Eur. J. Dermatol. 33, 368–382 (2023).

59. Haddad, H. K. et al. Hypothalamic opsin 3 suppresses MC4R signaling and potentiates Kir7.1 to promote food consumption. Proc. Natl. Acad. Sci. U. S. A. 122, e2403891122 (2025).

60. Mao, L. et al. Opsin 3 in keratinocytes mediates the light-induced exaggeration of atopic dermatitis. J. Allergy Clin. Immunol. 156, 980–992 (2025).

61. Nayak, G. et al. Adaptive Thermogenesis in Mice Is Enhanced by Opsin 3-Dependent Adipocyte Light Sensing. Cell Rep. 30, 672–686.e8 (2020).

62. Sato, M. et al. Cell-autonomous light sensitivity via Opsin3 regulates fuel utilization in brown adipocytes. PLoS Biol. 18, (2020).

63. Wu, A. D. et al. Opsin 3-Gαs Promotes Airway Smooth Muscle Relaxation Modulated by G Protein Receptor Kinase 2. Am. J. Respir. Cell Mol. Biol. 64, 59–68 (2021).

64. Yim, P. D. et al. Airway smooth muscle photorelaxation via opsin receptor activation. Am. J. Physiol. Lung Cell. Mol. Physiol. 316, L82–L93 (2019).

65. Koyanagi, M., Takada, E., Nagata, T., Tsukamoto, H. & Terakita, A. Homologs of vertebrate Opn3 potentially serve as a light sensor in nonphotoreceptive tissue. Proc. Natl. Acad. Sci. U. S. A. 110, 4998– 5003 (2013).

66. Sugihara, T., Nagata, T., Mason, B., Koyanagi, M. & Terakita, A. Absorption Characteristics of Vertebrate Non-Visual Opsin, Opn3. PLoS One 11, e0161215 (2016).

67. Wu, Q. et al. Opn3 Drives Blue-Light-Induced Reduction in Lipid Droplets and Antiviral Defense. Biomolecules 16, 109 (2026).

68. Jung, T., Höhn, A. & Grune, T. Lipofuscin: detection and quantification by microscopic techniques. Methods Mol. Biol. 594, 173–193 (2010).

69. Miyagi, M., Yokoyama, H., Shiraishi, H., Matsumoto, M. & Ishii, H. Simultaneous quantification of retinol, retinal, and retinoic acid isomers by high-performance liquid chromatography with a simple gradiation. J. Chromatogr. B Biomed. Sci. Appl. 757, 365–368 (2001).

70. Hughes, C. S. et al. Ultrasensitive proteome analysis using paramagnetic bead technology. Mol. Syst. Biol. 10, MSB145625-(2014).

71. Baker, C. P., Bruderer, R., Abbott, J., Arthur, J. S. C. & Brenes, A. J. Optimizing Spectronaut Search Parameters to Improve Data Quality with Minimal Proteome Coverage Reductions in DIA Analyses of Heterogeneous Samples. J. Proteome Res. 23, 1926–1936 (2024).

72. Ritchie, M. E. et al. limma powers differential expression analyses for RNA-sequencing and microarray studies. Nucleic Acids Res. 43, e47 (2015).

73. Perez-Riverol, Y. et al. The PRIDE database resources in 2022: a hub for mass spectrometry-based proteomics evidences. Nucleic Acids Res. 50, D543–D552 (2022).

74. Wang, M. et al. Assembling the Community-Scale Discoverable Human Proteome. Cell Syst. 7, 412 (2018).

