## Supplementary figure for "nnOPN3 mediates retinal-dependent lipofuscin accumulation and its loss sensitizes keratinocytes to blue-light-induced proteomic remodeling"

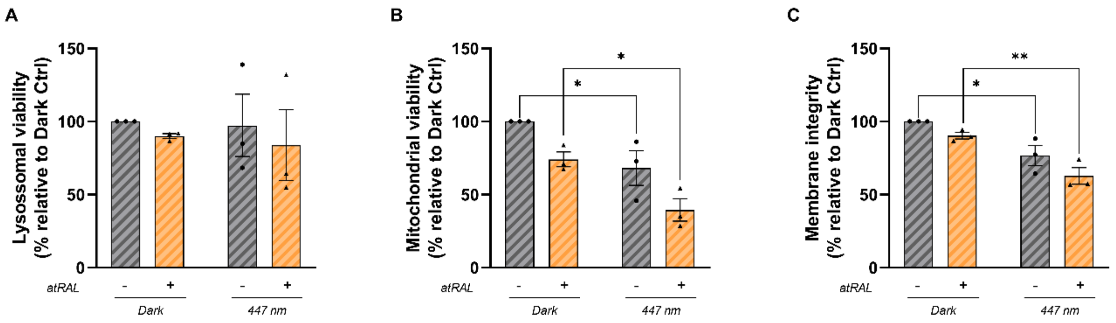


**Figure S1. Assessment of cell viability 48 h after atRAL photosensitization.** (A) Lysosomal viability assessed by the neutral red incorporation. (B) Mitochondrial viability assessed by the reduction of 3-(4,5-dimethylthiazol-2-yl)-2,5-diphenyltetrazolium bromide (MTT) to formazan. (C) Membrane integrity assessed by crystal violet staining. Experimental groups were Dark Control, Dark + atRAL, 447 nm Control, and 447 nm + atRAL. Each dot in the graphs represents one independent biological replicate. Data are presented as the mean ± SEM from at least three independent experiments. Statistical analysis was performed using two-way ANOVA followed by Bonferroni's multiple-comparisons test. ANOVA effects: Neutral red assay—interaction ($P$ = 0.9152); irradiation ($P$ = 0.7911); retinal treatment ($P$ = 0.4870). MTT assay—interaction ($P$ = 0.8561); irradiation ($P$ = 0.0023); retinal treatment ($P$ = 0.0067). Crystal violet assay—interaction ($P$ = 0.6648); irradiation ($P$ = 0.0006); retinal treatment ($P$ = 0.0344). $P < 0.05$.

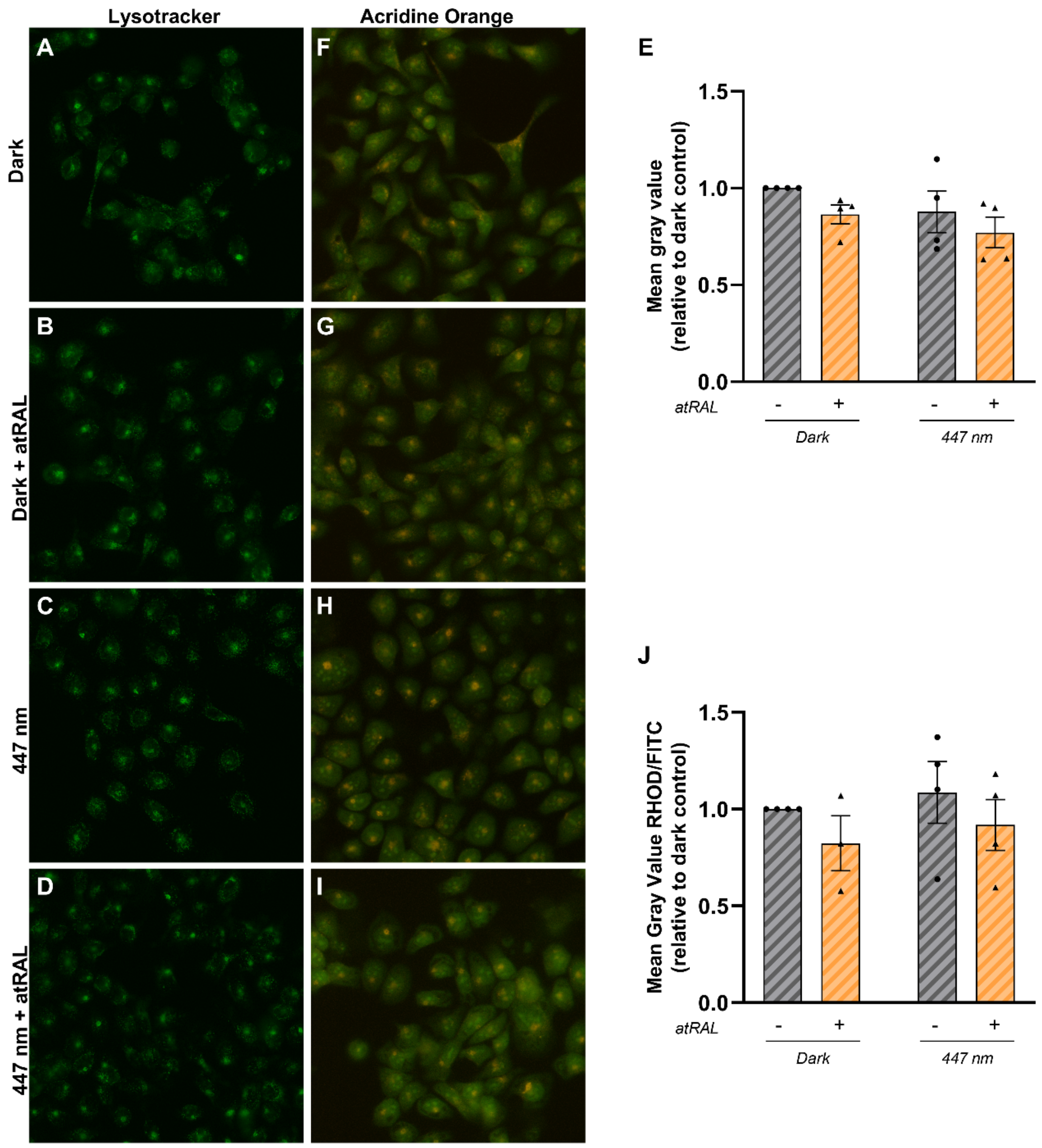


**Figure S2. Assessment of acidic vacuoles 3 h after atRAL photosensitization.** (A–D) Representative images of LysoTracker Deep Red staining in the different experimental groups. (E) Quantification of LysoTracker Deep Red fluorescence as mean gray value normalized to the Dark Control group. (F–I) Representative images of acridine orange fluorescence acquired in the FITC and Rhodamine (RHOD) channels. (J) Quantification of the acridine orange RHOD/FITC fluorescence ratio as mean gray value normalized to the Dark Control group. Experimental groups were: (A,F) Dark Control, (B,G) Dark + atRAL, (C,H) 447 nm Control, and (D,I) 447 nm + atRAL. A total of 50 cells per group were analyzed for each assay. Each dot in the graphs represents one independent biological replicate, with measurements obtained from the indicated number of cells per replicate. Data are presented as the mean ± SEM from at least three independent experiments. Statistical analysis was performed using two-way ANOVA followed by Bonferroni's multiple-comparisons test. **ANOVA effects:** LysoTracker Deep Red—interaction ($P = 0.8456$); irradiation ($P = 0.1553$); retinal treatment ($P = 0.1130$). Acridine orange RHOD/FITC ratio—interaction ($P = 0.9721$); irradiation ($P = 0.4831$); retinal treatment ($P = 0.1944$). $P < 0.05$.

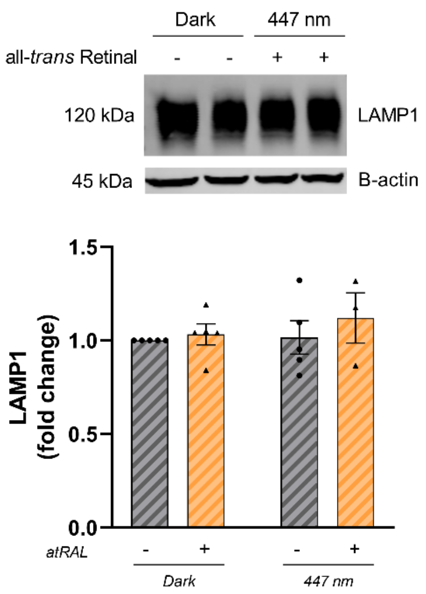


**Figure S3. Assessment of LAMP1 protein expression 24 h after atRAL photosensitization.** (A) Representative immunoblot of LAMP1 and β-actin. (B) Quantification of relative LAMP1 protein expression normalized to β-actin. Experimental groups were Dark Control, Dark + atRAL, 447 nm Control, and 447 nm + atRAL. Each dot in the graph represents one independent biological replicate. Data are presented as the mean ± SEM from at least three independent experiments. Statistical analysis was performed using two-way ANOVA followed by Bonferroni's multiple-comparisons test. ANOVA effects: interaction ($P$ = 0.6387); irradiation ($P$ = 0.4938); retinal treatment ($P$ = 0.3694). *$P$ < 0.05.

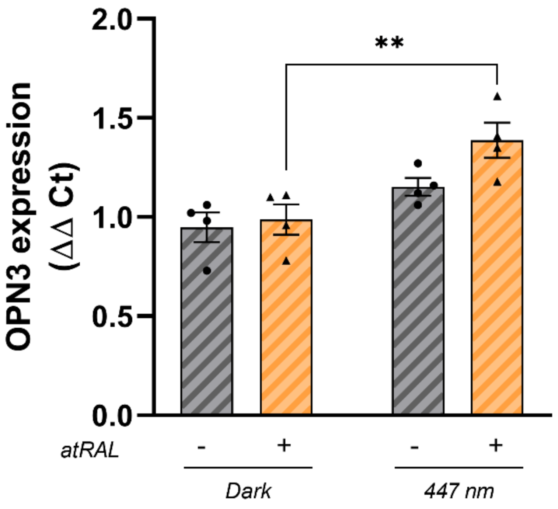


**Figure S4: Assessment of OPN3 gene expression 4 h after atRAL photosensitization.** Relative OPN3 mRNA expression measured by RT-qPCR and expressed as relative expression calculated using the ΔΔCt method. Experimental groups were Dark Control, Dark + atRAL, 447 nm Control, and 447 nm + atRAL. Each dot in the graph represents one independent biological replicate. Data are presented as the mean ± SEM from at least three independent experiments. Statistical analysis was performed using two-way ANOVA followed by Bonferroni's multiple-comparisons test. ANOVA effects: interaction ($P = 0.2064$); irradiation ($P = 0.0014$); retinal treatment ($P = 0.0840$). $^*P < 0.05$.

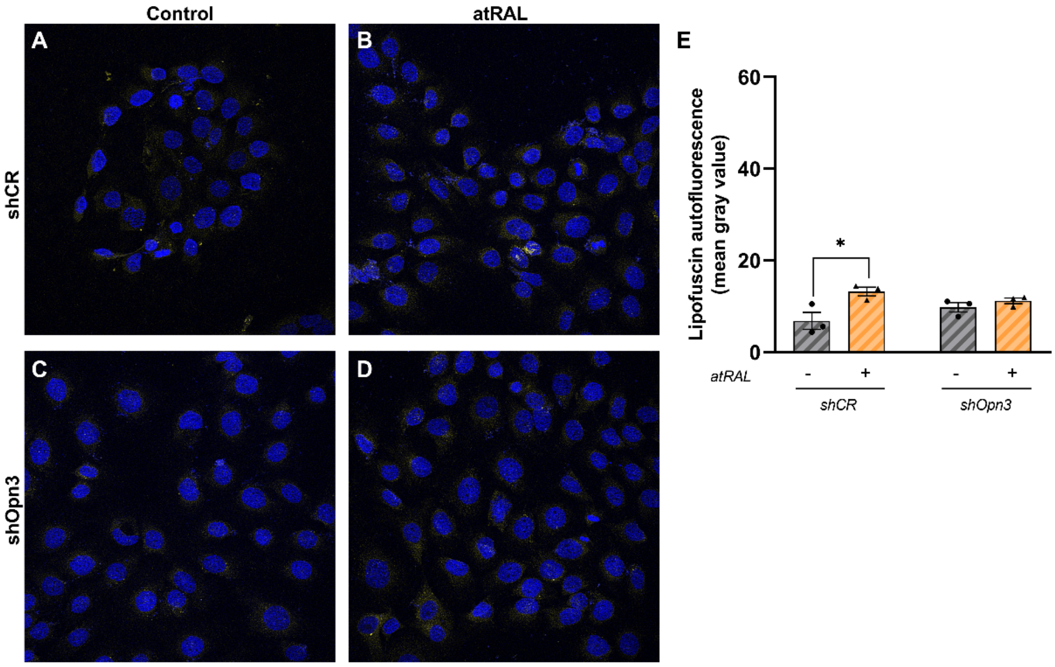


**Figure S5. Assessment of lipofuscin accumulation in OPN3-knockdown cells under non-irradiated conditions.** (A–D) Representative confocal images of lipofuscin autofluorescence (Ex. 488 nm/Em. 509–601 nm). (E) Quantification of lipofuscin autofluorescence as mean gray value. Experimental groups were: (A) Dark shCR Control, (B) Dark shCR + atRAL, (C) Dark shOpn3 Control, and (D) Dark shOpn3 + atRAL. A total of 50 cells per group were analyzed for lipofuscin autofluorescence. Each dot in the graph represents one independent biological replicate, with measurements obtained from the indicated number of cells per replicate. Data are presented as the mean ± SEM from at least three independent experiments. Statistical analysis was performed using two-way ANOVA followed by Bonferroni's multiple-comparisons test. ANOVA effects: interaction ($P$ = 0.0730); OPN3-knockdown ($P$ = 0.7087); retinal treatment ($P$ = 0.0122). *$P < 0.05$.

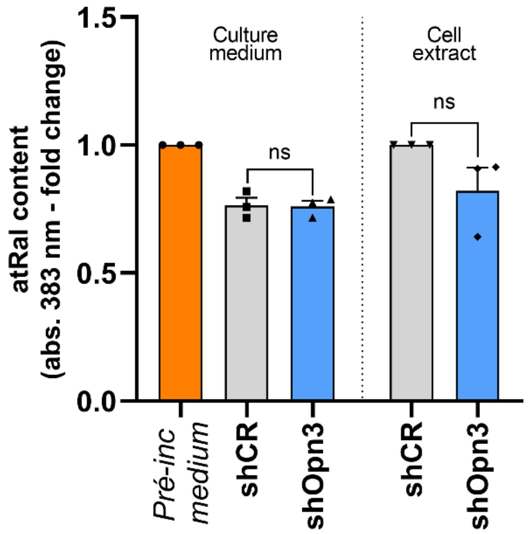


**Figure S6. Assessment of extracellular and intracellular atRAL levels in OPN3-knockdown cells following atRAL treatment.** The orange bar represents the atRAL concentration in the culture medium before incubation with the cells. Each dot in the graphs represents one independent biological replicate. Data are presented as the mean ± SEM from at least three independent experiments. Statistical analysis was performed using paired *t*-tests to compare extracellular atRAL levels between shCR and shOpn3 cells and intracellular atRAL levels between shSCR and shOPN3 cells. $P < 0.05$.

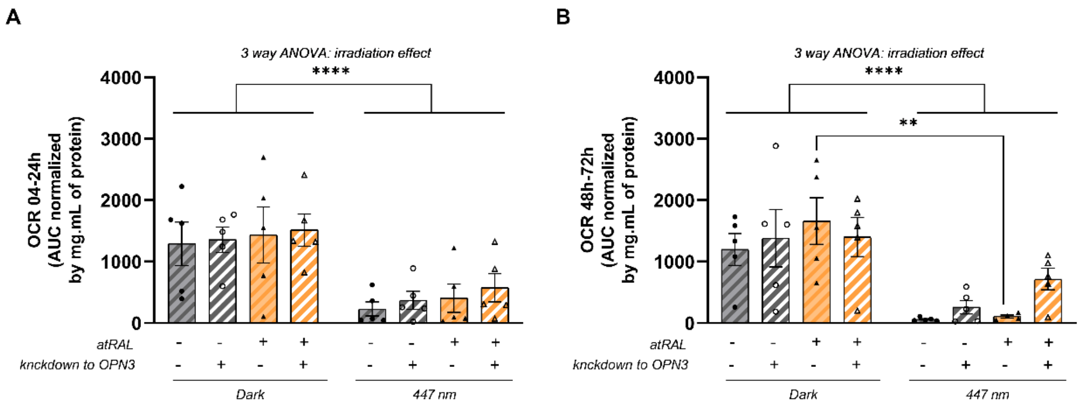


**Figure S7. Oxygen consumption in OPN3-knockdown cells following 447 nm irradiation and atRAL treatment.** (A) Quantification of oxygen consumption rate (OCR) as the area under the curve (AUC) from 0–24 h, normalized by total protein content. (B) Quantification of OCR as the AUC from 48–72 h, normalized by total protein content. Cells were treated with atRAL or vehicle and maintained in the dark or exposed to 447 nm blue light. Experimental groups consisted of shCR and shOPN3 cells in the presence or absence of atRAL under dark or 447 nm irradiation conditions. Each dot represents one independent biological replicate. Data are presented as the mean ± SEM from at least four independent experiments. Statistical analysis was performed using three-way ANOVA followed by Bonferroni's multiple-comparisons test. **ANOVA effects:** OCR 0 h – 24 h - irradiation ($P < 0.0001$); retinal treatment ($P = 0.3791$); knockdown ($P = 0.5539$); irradiation x retinal treatment ($P = 0.9175$); irradiation × knockdown ($P = 0.8311$); retinal treatment x knockdown ($P = 0.9562$); irradiation x retinal treatment x knockdown ($P = 0.9837$). OCR 0 h – 72 h - irradiation ($P < 0.0001$); retinal treatment ($P = 0.2168$); OPN3 knockdown ($P = 0.3589$); irradiation × retinal treatment ($P = 0.9774$); irradiation × OPN3 knockdown ($P = 0.2662$); retinal treatment × OPN3 knockdown ($P = 0.9671$); irradiation × retinal treatment × OPN3 knockdown ($P = 0.2837$). *$P < 0.05$.

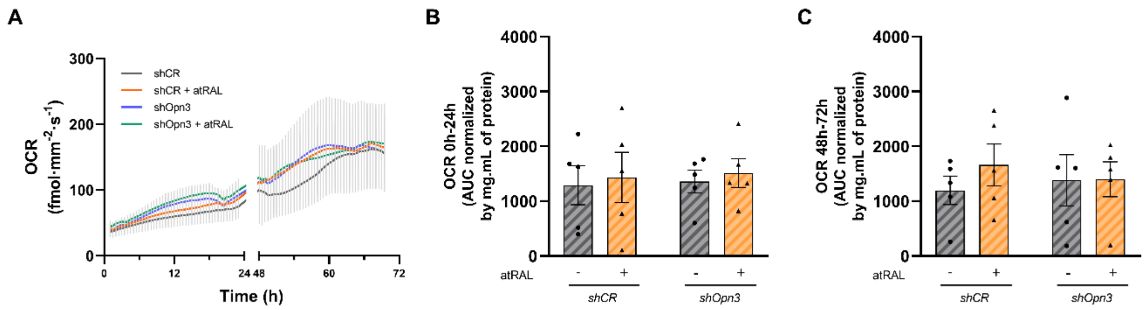


**Figure S8. Oxygen consumption in OPN3-knockdown cells following atRAL treatment under non-irradiated conditions.** (A) Oxygen consumption rates (OCRs) measured over 72 h in shCR and shOpn3 cells in the presence or absence of atRAL. (B) Quantification of OCR as the area under the curve (AUC) from 0–24 h, normalized by total protein content. (C) Quantification of OCR as the AUC from 48–72 h, normalized by total protein content. Each dot represents one independent biological replicate. Data are presented as the mean ± SEM from at least four independent experiments. Statistical analysis was performed using two-way ANOVA followed by Bonferroni's multiple-comparisons test. ANOVA effects: OCR 0 h – 24 h (B)—interaction ($P = 0.9844$); OPN3-knockdown ($P = 0.8290$); retinal treatment ($P = 0.6579$). OCR 0 h – 72 h (C)—interaction ($P = 0.5521$); OPN3-knockdown ($P = 0.9155$); retinal treatment ($P = 0.5181$). *$P < 0.05$.
